# Neurons in the human entorhinal cortex map abstract emotion space

**DOI:** 10.1101/2023.08.10.552884

**Authors:** Salman E. Qasim, Peter C. Reinacher, Armin Brandt, Andreas Schulze-Bonhage, Lukas Kunz

## Abstract

When animals move through space, neurons in their entorhinal cortex activate periodically at multiple locations to form a map of the spatial environment. These grid cells may also map non-physical, conceptual spaces to support various other complex behaviors. Here, using intracranial recordings in neurosurgical patients performing an emotional memory task, we find that neurons in the human medial temporal lobe activate in a grid-like pattern across a two-dimensional feature space characterized by emotional valence and arousal. These neurons were different from cells tuned solely to valence or arousal, were preferentially located in the entorhinal cortex, and exhibited theta-phase locking. Our observation of grid-like neuronal activity during emotional processing in humans supports the idea that the neural structure of cognitive maps generalizes beyond spatial navigation.

## Introduction

To perform spatial navigation, animals and humans appear to develop cognitive maps of their environment^1–3^. Cognitive maps encode the relationships between spatial locations and thus help animals and humans to orient themselves relative to their surroundings, make predictions about where future paths will lead them, and infer new routes through space^4,5^. These functions of knowledge organization, prediction, and inference are also critical for many non-spatial behaviors^1,6–8^. The use of cognitive maps may thus generalize beyond spatial navigation and facilitate cognitive computations across a variety of behavioral tasks in which information is distributed along different feature dimensions^9,10^.

Electrophysiological recordings in rodents have identified neurons in the medial temporal lobe (MTL) that are tuned to spatial locations in a way that could form the neural substrate of cognitive maps^11,12^. Among these neurons are entorhinal grid cells that activate periodically at multiple, hexagonally arranged locations when rodents and other species move through two-dimensional space^13–15^. They form a population code of the subject’s position^16^ and may support navigational processes such as the computation of vectors to goals^17–19^. Combining the discovery of grid cells with the idea that cognitive maps are relevant beyond spatial navigation, researchers have recently started to demonstrate the presence of grid-cell spiking when animals explore visual and auditory space^20,21^. Human neuroimaging studies furthermore identified a putative proxy for population-level grid-cell activity, and studied this population signal when humans navigated through various sensory or abstract conceptual spaces^22–29^.

Whether grid cells might also be involved in emotional processing has remained unknown, however. Hence, building on the idea of a generalized role for grid cells across different behavioral domains^9,10^, we tested whether neurons in the human medial temporal lobe exhibit grid-like activity (i.e., periodic spiking reminiscent of grid-cell firing) to map an abstract, conceptual space spanned by the emotional features of valence and arousal. Valence and arousal are considered orthogonal basic dimensions of emotion^30,31^ and capture how pleasant (negative to positive) and how intense (relaxing to engaging) emotional experiences are, respectively^32,33^. We reasoned that the observation of grid-like activity in such a two-dimensional emotion space could indicate a new behavioral function for grid-like neuronal activity in the brain and would provide new insights into the neural mechanisms of emotional cognition in humans^34,35^.

## Results

### Participants encode and recall images in a two-dimensional emotion space

To investigate whether human neurons exhibit grid-like spiking in an emotional space with valence and arousal as the two dimensions, we recorded single-neuron activity in epilepsy patients as they performed a memory task with emotional images (13 patients; 14 experimental sessions). During every trial of this memory task, participants encoded a series of 12 images and, after a distractor period, freely recalled the images in any order (Fig. 1A). Participants completed 20 trials per session, encoding a total of 240 images. They were able to freely recall 47.7 ± 11.3% (mean ± standard deviation) of these images on average (Fig. 1B), indicating that they were attentive to the content of the images.

**Figure 1:**
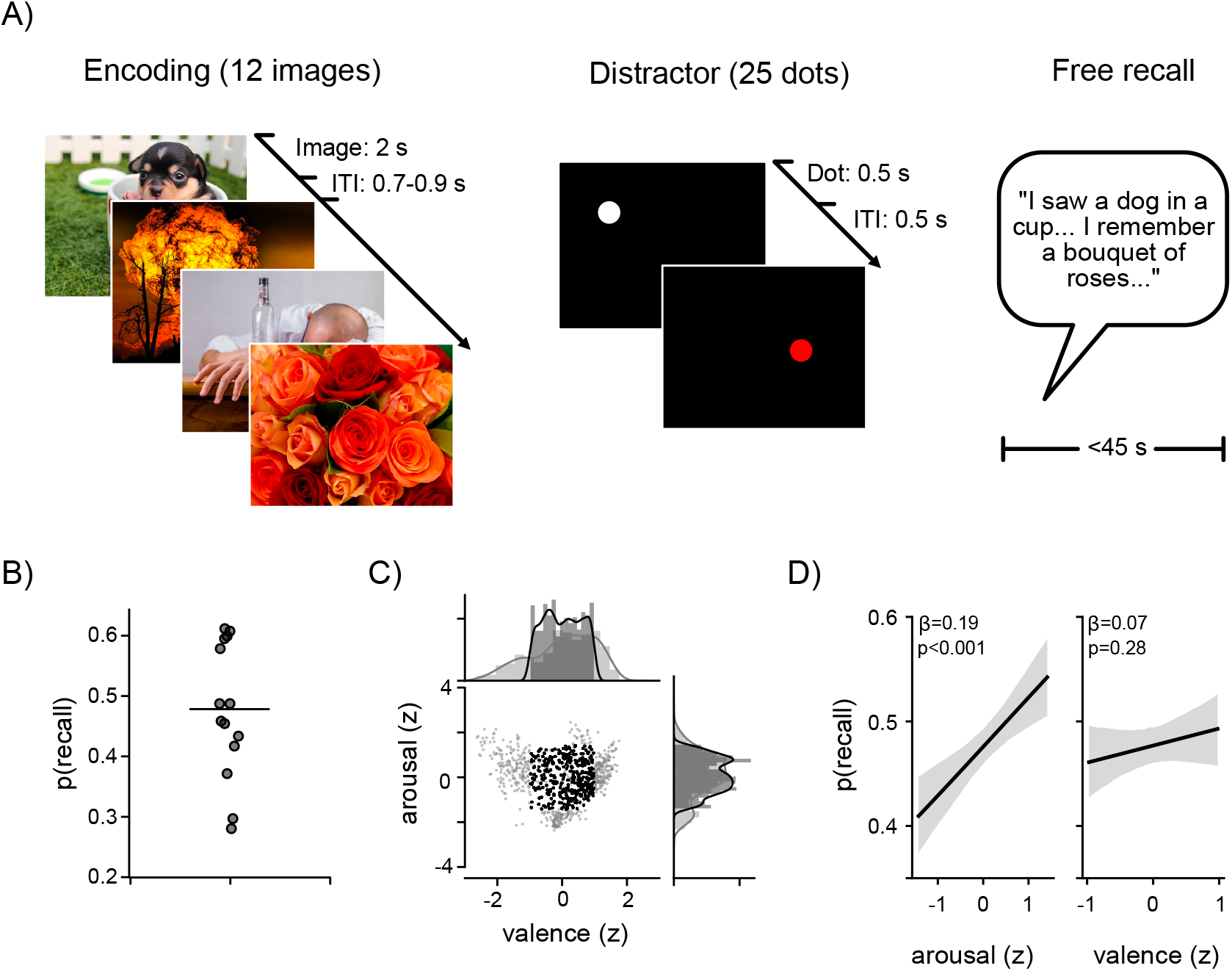
Emotional memory task. (**A**) Each task session consisted of 20 trials, each of which comprised an encoding period, a distractor period, and a free recall period. During the encoding period, participants viewed 12 different images (2-s duration per image; 0.8-s inter-image interval [ITI], jittered). During the distractor period, they viewed white and red dots at different locations on the laptop screen (0.5-s duration per dot, slightly jittered; 0.5-s inter-dot interval, slightly jittered) and were asked to press a button whenever a red dot appeared. During the free recall period, participants recalled the images from the preceding encoding period and described them aloud (recorded by a microphone). (**B**) Probability of recall for all participants (one dot per participant). Horizontal line denotes the mean recall probability. (**C**) Joint distribution of z-scored arousal and valence ratings for all images from the OASIS database (gray; *n* = 900) and the subset of images selected for this study (black; *n* = 377). (**D**) Logistic model fits of the probability of recall as a function of z-scored arousal (left) and valence (right). Shading denotes 95th percentile of bootstrapped model fits. Beta coefficient is indicated above model fit, along with the *P*-value for the coefficient *t*-statistic.

All images presented during the task were drawn from the Open Affective Standardized Image Set (OASIS), which is a large database of images with high-reliability valence and arousal ratings from an independent group of participants^36^. This allowed us to assign each image a specific location in emotional valence–arousal space and to later investigate neural spiking as a function of location in this space. We specifically used a subset of the OASIS images to sample a relatively uniform, square subpart of the valence–arousal space, motivated by grid-cell studies in rodents in which animals move through square boxes (Fig. 1C)^13^. Similar to previous work in epilepsy patients^37^, images drawn from this part of the emotional space were more likely to be recalled if they had higher arousal ratings, whereas the effects of valence on recall were more variable (Fig. 1D).

### Neuronal spiking is modulated by arousal and valence

We recorded neuronal activity from microwire bundles^38^ implanted in the entorhinal cortex, amygdala, hippocampus, and parahippocampal cortex (Figs. 2A; S1) as participants performed the emotional memory task. We identified 461 putative single neurons in these regions (Fig. S2; see *Methods*) and, as a first step, examined the neurons’ overall activity levels during the encoding periods of the task. Averaging across all images, we observed that neurons in all four brain regions generally increased their spiking during encoding (Fig. 2B), with parahippocampal neurons showing the strongest and earliest firing-rate increase, in line with previous work^39^.

**Figure 2:**
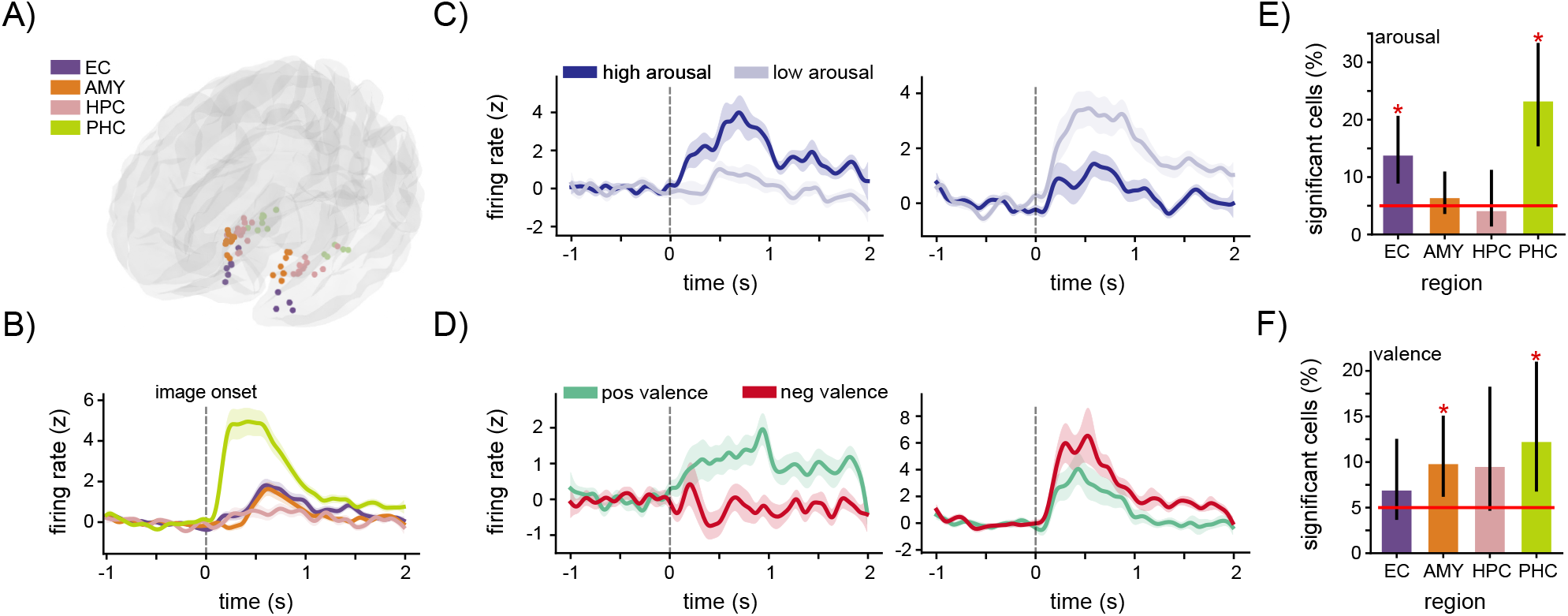
Neuronal spiking is modulated by the emotional features of images during memory encoding. (**A**) Template brain depicting the putative location of all microwire bundles, with color indicating the brain region of each bundle. (**B**) Mean firing rate (z-scored relative to the 1-s baseline and 2-s encoding period) of neurons in each region time-locked to image presentation during encoding (dotted line, image onset). Parahippocampal neurons responded most quickly and strongly to image presentation. Shaded areas denote the standard error. (**C**) Neurons were tuned to arousal during emotional memory encoding. Left: mean firing rate (z-scored) of neurons (*n* = 16) that exhibited a positive relationship between arousal and firing rate time-locked to image presentation during encoding (dotted line, image onset). Right: mean firing rate (z-scored) of neurons (*n* = 35) that exhibited a negative relationship between arousal and firing rate time-locked to image presentation during encoding (dotted line, image onset). Shaded areas denote the standard error. (**D**) Neurons were tuned to valence during emotional memory encoding. Left: mean firing rate (z-scored) of neurons (*n* = 24) that exhibited a positive relationship between valence and firing rate time-locked to image presentation during encoding (dotted line, image onset). Right: mean firing rate (z-scored) of neurons (*n* = 19) that exhibited a negative relationship between valence and firing rate time-locked to image presentation during encoding (dotted line, image onset). Shaded areas denote the standard error. (**E**) Percentage of putative single units in each region that exhibited significant arousal tuning. Vertical lines denote 95% binomial confidence intervals. Red line denotes chance level (5%). Asterisks denote significant percentages (binomial tests, *P*_corr._ < 0.05, Bonferroni corrected for four tests). (**F**) Percentage of putative single units in each region that exhibited significant valence tuning. Vertical lines denote 95% binomial confidence intervals. Red line denotes chance level (5%). Asterisks denote significant percentages (binomial tests, *P*_corr._ < 0.05). AMY, amygdala; EC, entorhinal cortex; HPC, hippocampus; PHC, parahippocampal cortex.

We next tested whether arousal and valence ratings modulated the tuning of individual neurons in the medial temporal lobe. Out of the 461 neurons, 51 neurons (11.1%) significantly altered their firing rates as a function of image arousal, either increasing (Figs. S3A; 2C, left) or decreasing (Figs. S3A; 2C, right) their activity when participants encoded images with higher arousal ratings (tertile split). Similarly, 43 neurons (9.3%) significantly altered their firing rates as a function of image valence, either increasing their activity during the encoding of positive images (Figs. S3B; 2D, left), or during the encoding of negative images (Figs. S3B; 2D, right). We observed significant percentages of arousal-tuned neurons in entorhinal and parahippocampal cortices (binomial tests, all *P*_corr._ < 0.004, Bonferroni corrected for four tests), and of valence-tuned neurons in amygdala and parahippocampal cortex (binomial tests, all *P*_corr._ < 0.033). An insignificant number of neurons (5/461) exhibited conjunctive coding for both arousal and valence. Adding to prior work demonstrating how emotional features modulate spiking in the subthalamic nucleus^40^, these results demonstrate that valence and arousal ratings modulate the spiking of individual neurons in the medial temporal lobe and thus provide new insights into how basic emotional features are separately represented in the human brain.

### Periodic spiking in the entorhinal cortex maps 2D emotion space similar to grid cells

To assess the possibility that neurons in the human brain form a cognitive map of emotion space, we next examined grid-like tuning in the two-dimensional valence–arousal coordinate system. For each neuron, we thus computed a two-dimensional firing-rate map in emotion space (Figs. 3A; S4; see *Methods*) and observed cells that exhibited multi-peaked spiking in this space (Figs. 3B; S5). To systematically characterize the spatial tuning of these neurons, we computed the grid score and spatial information of each neuron’s firing-rate map in emotion space, analogous to the metrics used to measure the activity of grid and place cells in rodents navigating physical space^41,42^. We estimated grid scores using the spatial autocorrelogram of spiking in emotion space, which showed evidence for regularly spaced peaks at multiples of 60^◦^, and compared these grid scores to surrogate grid scores from the same cells to determine significant grid-like spiking (Figs. 3A; S6; see *Methods*). Across the entire population of 461 neurons, we identified 28 neurons that exhibited significant grid-like spiking (Figs. 3B; S5).

**Figure 3:**
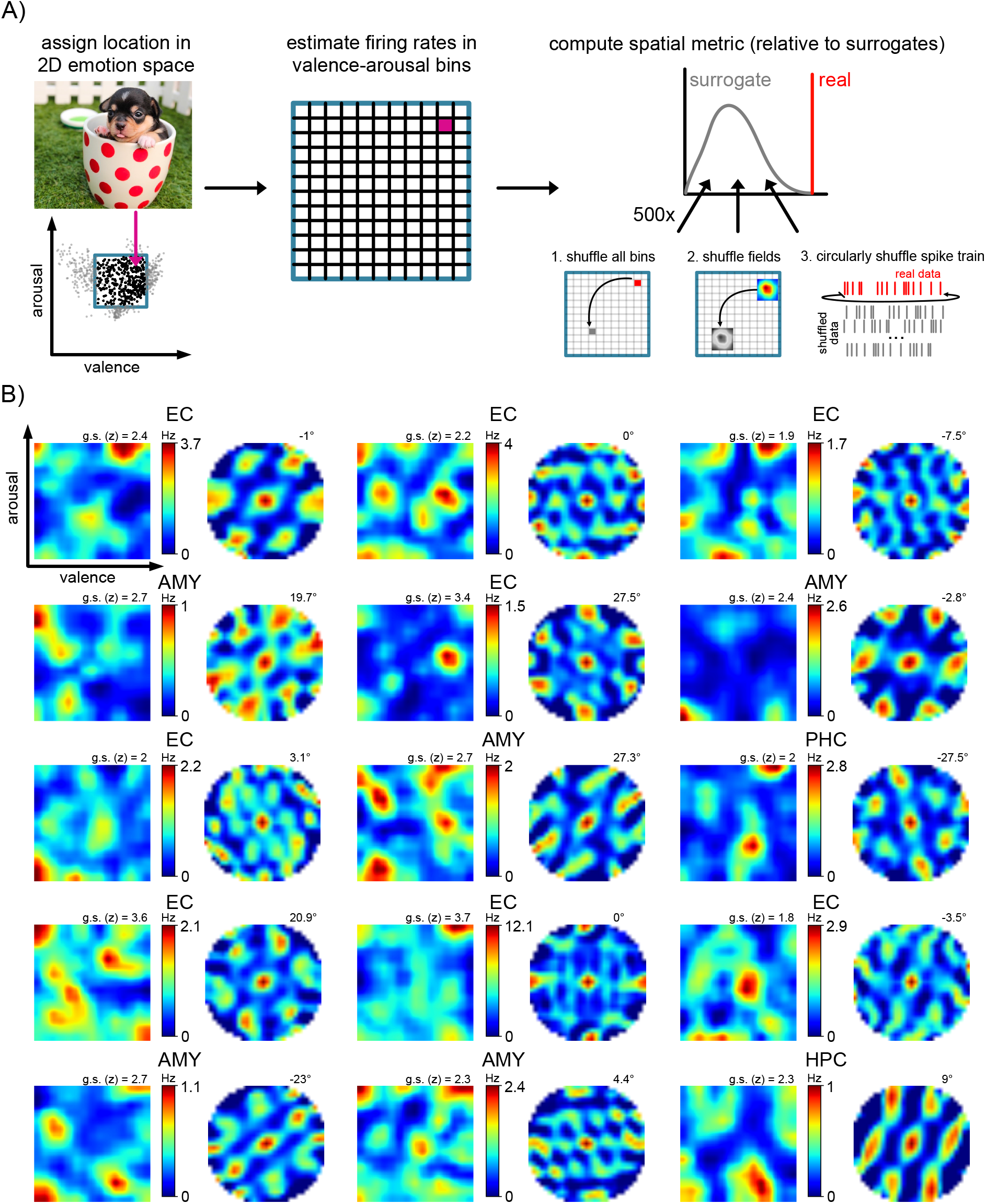
Measuring neuronal activity in emotion space reveals grid-like spiking. (**A**) Schematic of analysis approach. We assigned images a position in a two-dimensional emotion space based on their valence and arousal ratings. We binned neuronal spiking in this emotion space, and computed spatial metrics such as grid scores. We generated surrogate-corrected spatial metric scores using empirically-generated surrogate null distributions (see *Methods*). (**B**) Examples of grid-like spiking activity in emotion space. For each neuron, firing-rate maps in emotion space are depicted on the left, with colorbar indicating minimum and maximum firing rates. Warm colors denote higher firing rates, cool colors denote lower firing rates. Spatial autocorrelogram is depicted on the right, with the grid-orientation angle stated at the top right. Each neurons’ brain region is indicated above. AMY, amygdala; EC, entorhinal cortex; HPC, hippocampus; PHC, parahippocampal cortex; g.s., grid score.

Alternative geometric arrangements (e.g., 90^◦^ angular spacing) did not explain the activity of these grid-like neurons (Fig. 4A). Additionally, grid-like firing was not explained by tuning to valence or arousal alone (Fig. 4B). In support of the temporal stability of their tuning, grid-like neurons exhibited significantly correlated spatial firing across the first and second halves of the task (one-sample *t*-test, *t*_27_ = 3.2, *P* = 0.003), which was also stronger than in other, non-grid-like neurons (two-sample *t-*test, *t*_459_ = 2.4, *P* = 0.015; Fig. 4C). Of the 28 grid-like neurons we identified, only the entorhinal cortex exhibited a significant proportion of such cells (*n* = 14; binomial test, *P*_corr._ = 0.029, Bonferroni corrected for four tests; Fig. 4D), in line with the localization of grid cells to the entorhinal cortex in rodents navigating physical space^13^. These neurons were identified in 7 of 10 participants with electrodes in the entorhinal cortex (Table S1).

**Figure 4:**
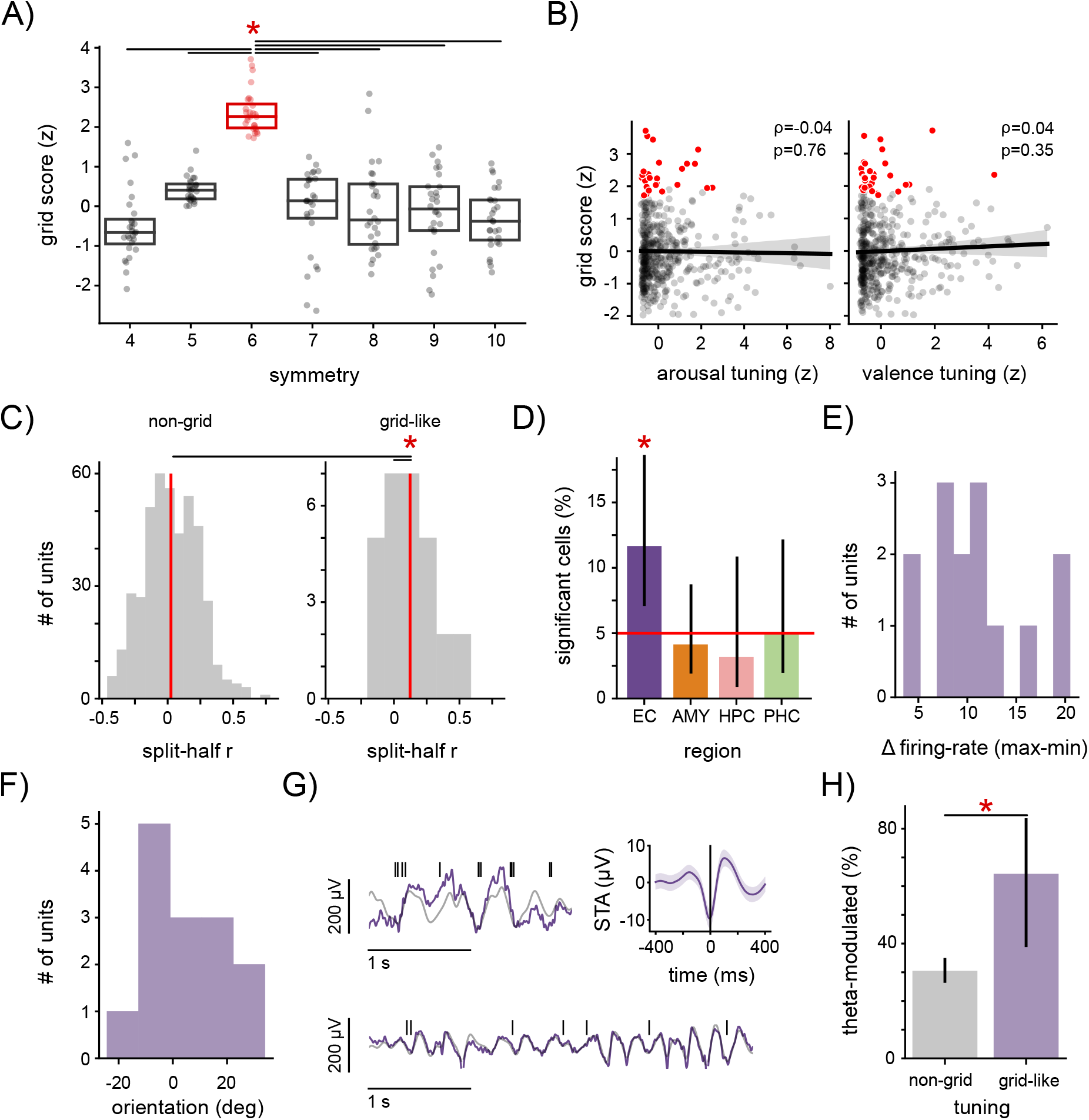
Neurons with grid-like spiking are located in the entorhinal cortex and are theta modulated. (**A**) Grid-like spiking in emotion space shows 6-fold symmetry rather than other symmetries. Grid score for each grid-like neuron (circle) as a function of various symmetries. Horizontal lines denote paired comparisons between 6-fold symmetry and all other symmetries. Asterisk denotes significance of all paired comparisons. (**B**) Non-significant correlations between arousal/valence tuning and emotion grid scores. Lines denote linear regression fits. Shaded areas denote bootstrapped 95% confidence intervals. Red dots denote grid-like neurons across all brain regions. (**C**) Split-half correlations for non-grid-like (left) and grid-like neurons (right), quantifying the temporal stability of their spiking in emotion space. Vertical, red lines denote the mean of each distribution. Asterisk indicates a significant one-sample *t*-test against *r* = 0 for the population of grid-like neurons, and a significant two-sample *t*-test between the two cell populations (both *P* < 0.05). (**D**) Percentage of putative single units in each region that exhibited significant grid scores. Vertical lines denote 95% binomial confidence intervals. Red line denotes chance level (5%). Asterisks denote significant percentages (binomial test, *P*_corr._ < 0.05, Bonferroni corrected for four tests). (**E**) Range of binned firing rates in the firing-rate maps for grid-like neurons in the entorhinal cortex. (**F**) Range of grid orientations for grid-like neurons in the entorhinal cortex. (**G**) Left: examples of local-field potential (LFP) activity (purple), with 2–10 Hz phase estimate overlaid (gray) and corresponding spiking activity depicted above for two example grid-like cells. Right: spike-triggered average (STA) across all entorhinal grid-like neurons. (**H**) Percentage of significantly theta-modulated neurons in the entorhinal cortex, split into grid-like neurons (purple) and non-grid-like neurons (gray). Vertical lines denote 95% binomial confidence intervals. Asterisk denotes significant difference in percentages (χ^2^ test of proportions, *P* < 0.05). #, number.

Grid-like neurons in the entorhinal cortex exhibited a larger range of firing rates in emotion space than non-grid-like neurons (two-sample *t*-test, *t*_445_ = 2.1, *P* = 0.03; Fig. 4E), and exhibited a range of grid orientations consistent with prior work in rodents^43^ (Fig. 4F). Grid cells in rodents show significant theta modulation^44^, so we next tested whether grid-like spiking in emotion space was locked to the theta oscillation. Because human theta is more transient and non-stationary than in rodents^45,46^, we computed theta modulation using spike-phase autocorrelations within a 2–10 Hz frequency range^47,48^. A significant number of entorhinal grid-like neurons (9/14; binomial test, *P* < 0.001) exhibited clear phase-locking to the theta oscillation, with neuronal spiking aligning to the theta cycle even if the oscillation was irregular (Fig. 4G, left). Grid-like neurons were specifically locked to the trough of theta, in line with findings in rodent and monkey grid cells^20,44,49^ (Fig. 4G, right). Theta-phase-locked neurons were significantly more prevalent among grid-like neurons in the entorhinal cortex than non-grid-like neurons (χ^2^-test for proportions, χ^2^ = 7.2, *P* = 0.007; Fig. 4H).

### Control analyses of grid-like spiking during emotional processing

We identified grid-like activity in emotion space using surrogate-corrected grid scores (Figs. 3; S6; see *Methods*). We performed further analyses to rule out that this phenomenon could potentially be explained by our participants’ behavioral sampling of emotion space. In two participants, we observed that the occupancy maps in emotion space showed grid-like patterns by chance, but on the whole occupancy gridness did not predict cellular grid-like tuning (mixed-effects linear model, *P* = 0.86; Fig. S7).

Furthermore, to identify if behavioral sampling could lead to false-positive grid cells, we applied our analysis to a “ground truth” dataset of grid and non-grid cells from rodents moving through physical space (Fig. S8A)^41^ where we masked the firing-rate maps with our participants’ occupancy maps in emotion space (Fig. S8B). Grid scores for the masked rodent data were highly correlated with grid scores from the raw, unmasked data (Pearson’s correlations, all ρ > 0.81, all *P* < 0.001; Fig. S8C). Furthermore, when used to classify cells as grid cells, the masked rodent data did not result in elevated false-positive rates (area under the ROC curve [AUC] = 0.87; Fig. S8D), demonstrating that inhomogeneous or incomplete sampling of emotion space did not bias our analysis of grid-like tuning toward higher numbers of false positives. We also examined the effect of different resolutions when binning the emotional space and found that different bin counts led to highly similar results (Fig. S8C). As a resolution of 15 x 15 bins resulted in the closest relationship between grid scores from the masked and unmasked firing-rate maps in rodents, we used this bin count to analyze grid-like tuning in our data (see above). Overall, behavioral sampling did not relevantly affect grid-like tuning in emotion space.

We next assessed whether grid-like activity might be explained by other cognitive demands of the task. Specifically, the semantic category of an image contributed significantly to the likelihood of subsequent recall (Fig. S9A) and many neurons in this task were selectively tuned to a particular image category (responding preferentially to objects, scenes, persons, or animals; Fig. S9, B–D). We found, however, that grid-like firing was not correlated with the cells’ tuning to category (Fig. S9E). Hence, because valence, arousal, and category tuning did not appear to be associated with grid scores (Figs. 4B; S9E), it indicates that grid-like tuning in emotion space was not simply reducible to other forms of neural tuning to behaviorally relevant information.

Finally, we tested whether the grid-like neurons we identified in emotion space also exhibited grid-like spiking in visual space, as visually-evoked grid-like neural representations have been identified in studies using monkey electrophysiology^20^ and human neuroimaging^23,24^. We thus applied our analysis of grid-like tuning to the neural data from the distractor period of the task, when participants viewed dots appearing at various positions on the screen. Because participants attended to the dots, we considered the position of the dots as a proxy for visual fixation position. We computed the firing-rate map of each neuron’s spiking activity in visual space and computed surrogate-corrected grid scores for each neuron in this space (Fig. S10, A–B). Though we identified examples of grid-like activity in visual space (Fig. S10C), we did not identify a significant proportion of such cells in any region (Fig. S10D). Furthermore, visual grid scores were not correlated with emotional grid scores, suggesting that grid-like activity in visual space did not account for grid-like activity in emotional space (Fig. S10E).

## Discussion

Our results are consistent with the idea that grid cells are a neural substrate of cognitive maps that structure knowledge in spatial and non-spatial feature dimensions^9,10^. Humans are thought to use such knowledge to guide learning and inference^50^ and to generalize their behavior across tasks^51^. Here, using single-neuron recordings in humans encoding emotional images, we found that neurons in the human brain show periodic, multi-peaked spiking when their firing rates were quantified as a function of the images’ position in a two-dimensional valence–arousal emotion space. Cells with such grid-like tuning were most prevalent in the entorhinal cortex; showed a preference for hexagonal symmetry; could not be explained by pure valence, arousal, or category tuning; and showed theta-phase locking. These findings suggest that basic emotional features can serve as dimensions for cognitive maps that involve the activity of entorhinal grid cells.

A variety of previous studies in rodents described the characteristic hexagonal firing pattern of grid cells when the animals moved through two-dimensional physical space^52^. As a population, grid cells represent a subject’s exact position in space^16^. Grid-like spiking in spaces defined by basic emotional dimensions may thus provide participants with knowledge about their current emotional state. Theoretical and computational studies furthermore indicate that grid cells are useful for computing vectors to goals^17,18^, suggesting that grid cells help participants to infer their future locations in space and to plan trajectories towards these locations^53^. In the context of emotional processing, such inferences would allow participants to estimate the emotional consequences of their actions, which may support their decision-making processes^54^. As emotions are critical components of interpersonal communication, the ability of predicting emotional states via grid cells could also be relevant for social cognition^55^.

Our findings build upon a model that decomposes complex emotional states into basic, orthogonal emotional features that serve as continuous emotional dimensions^31,32,56^. Other theories propose a limited, discrete set of basic emotions, such as anger and fear, that are qualitatively different from each other^57^. Together with prior neuroimaging studies on valence and arousal^58–61^, the neural findings of this study are consistent with the former, dimensional model of emotion, but they cannot provide decisive evidence in favor of or against either theory. Models of discretely partitioned emotional states may still account for the formation of grid-like representations, as certain computational models posit that grid-cell activity emerges via the clustering of concepts during learning^62^. Furthermore, it may well be that the brain contains both neural codes for representing continuous emotional dimensions (for example in the entorhinal cortex) and for representing particular, discrete emotional states (potentially in the amygdala or hippocampus), but this idea remains an open question.

By demonstrating that human entorhinal neurons show grid-like tuning in a two-dimensional valence–arousal space, our findings provide a link between rodent electrophysiology studies of spatial navigation, human neuroimaging studies of cognition and emotion, and theoretical and computational models linking these different lines of research under the auspice of a cognitive mapping framework. Future work may investigate neuronal spiking in larger emotional spaces as extending the size of the spatial enclosure helped identify the firing pattern of grid cells in early rodent grid-cell studies by recording more firing fields^13,63^. High-density recordings from the human entorhinal cortex^64,65^ may determine whether the neurons we identified exhibit similar modularity as grid cells in rodent entorhinal cortex, with comparisons of grid scale and orientation across multiple neurons from the same recording site^43^. By asking participants to provide their own valence and arousal rating of each image, future studies may furthermore test whether grid-like spiking in two-dimensional valence–arousal space is stronger when considering subjective rather than crowdsourced (objective) ratings^66^. They may also add a third emotional dimension^67^, in analogy to the investigation of grid cells in three-dimensional physical space^68,69^. Finally, future studies may investigate if alternative, non-grid representations (such as graph-like representations) better capture the activity of neurons in abstract concept space^70,71^. Together, this research may increase our understanding of the cellular basis of emotions and the functional role of grid cells.

## Methods

### Human participants

We recorded data from 15 patients undergoing invasive intracranial EEG monitoring in the course of their treatment for drug-resistant epilepsy at the Freiburg Epilepsy Center, Freiburg im Breisgau, Germany. The location and number of implanted electrodes varied between patients and was determined solely as as a function of clinical needs. Two participants were excluded due to a lack of verbal recall responses during the task, leaving 13 participants (9 female; age = 37.8 ± 14 years [mean ± standard deviation]; Table S1). One participant completed two sessions, leading to a total number of 14 experimental sessions. The study conformed to the guidelines of the ethics committee of the University Hospital Freiburg, Freiburg im Breisgau, Germany, and all patients provided written informed consent.

### Neurophysiological recordings

Electrophysiological data for the detection of putative single units were recorded using microwire electrodes protruding from the tip of penetrating depth electrodes^38^ (Ad-Tech Medical Instruments, Racine, WI, USA) at 30 kHz using a NeuroPort system (Blackrock Microsystems, Salt Lake City, UT, USA). Each Behnke-Fried microelectrode bundle consists of 8 platinum–iridium microelectrodes with a diameter of 40 µm, with a 9th, uninsulated microelectrode serving as a reference. We recorded from the amygdala (AMY), hippocampus (HPC), parahippocampal cortex (PHC) and entorhinal cortex (EC). We assigned the microelectrode bundles to these brain regions by visually inspecting the trajectories of the implanted depth electrodes on post-implantation MRI scans.

### Behavioral task

During their hospital stay, participants sat in their hospital bed and performed a memory task while neural data was recorded. The task consisted of 20 trials, each of which comprised an encoding period, a distractor period, and a free recall period.

During the encoding period, 12 different images were presented to the subject. The images were drawn from the Open Affective Standardized Image Set (OASIS), in which they were previously rated with regard to valence and arousal by an independent group of healthy participants^36^. We selected images so that they spanned a square subpart of the valence–arousal space of the database, using images with a valence rating of 4.33 ± 1.2, and an arousal rating of 3.67 ± 1.2. Each image was presented for a duration of 2 seconds, with an inter-image interval of 0.8 seconds (randomly jittered between 0.7 and 0.9 seconds). Participants were asked to encode these images to recall them during the subsequent retrieval period. During the distractor period, the participant viewed white and red dots at different locations on the laptop screen (randomly drawn from a square area of size 810 × 810 pixels, centered on the laptop screen, which was of size 1920 × 1080 pixels). Each dot was shown for a duration of 0.5 seconds (slightly jittered), with an inter-dot interval of 0.5 seconds (slightly jittered). The participants were asked to press a button whenever a red dot appeared. For the first participant, 20 dots were shown during each distractor period; for all other participants, 25 dots were shown during each distractor period. Participants responded accurately to the dots on 77 ± 12% of the trials (mean ± standard deviation), supporting the idea that they visually attended to the dots and their locations on the screen. During the free recall period, the participants then freely recalled the images from the preceding encoding period and described them aloud (recorded by a microphone). Each free recall period had a maximum duration of 45 seconds, but participants were able to terminate a recall period ahead of time by pressing the space bar. No feedback about recall accuracy was provided during the task.

Participants did not undergo training during the task to develop a (subconscious) understanding of the emotional space, as was done in previous studies on abstract conceptual spaces^22^. Participants thus encountered the paradigm simply as a memory task (experiencing valence and arousal as intrinsic features of the images) and were not aware that the images could be organized in a two-dimensional emotion space.

### Data processing and analysis tools

All pre-processing and analyses were performed using Python. Statistical analyses were carried out primarily using the scipy^72^ and statsmodels^73^ toolboxes and custom Python functions. All custom Python scripts will be publicly available on GitHub upon publication. All figures were made using the Matplotlib^74^ and Seaborn libraries^75^.

### Behavioral analysis

We annotated and scored audio recordings from the free recall period using the ELAN software^76^. For each recall, we identified the start time, duration, and verbal response of the participant. Based on the participant’s verbal response, we identified the image from the encoding list the response referred to. In rare cases, a participant’s description of a remembered image referred to more than one image from encoding. We coded these ambiguous recalls to serve as a recall for all applicable images in the encoding list. Recall probability was computed as the number of correct recalls divided by the total number of images encoded. To investigate the relationship between emotional characteristics of the images and memory, we extracted arousal and valence ratings from the OASIS database, z-scored these ratings relative to all of the images in the database, and assigned them to each stimulus. We then fit a logistic regression model to assess the contribution of valence and arousal ratings to recall probability (Fig. 1D). We aligned the behavioral data with the neural data using visual triggers detected by a phototransistor attached to the screen of the task computer and recorded as an analog signal by the NeuroPort recording system.

### Preprocessing of neural data

We utilized Combinato^77^ to identify putative single neurons from the recorded electrophysiology data. Manual sorting identified single-versus multi-unit activity versus noise on the basis of previously determined criteria including (but not limited to) waveform shape and inter-spike interval (ISI) distribution^78,79^. The local field potential (LFP) for each neuron was recorded from the local microelectrodes and was downsampled to 250 Hz for spectral analysis. We identified 464 putative single-units, 3 of which were excluded due to low firing rates (< 0.1 Hz). Putative multi-units were excluded from all analyses.

### Identifying tuning to arousal, valence, and category

For each neuron, we fit a linear model to the relationship between trial-wise firing rate during the 2-s encoding period, and arousal, valence, and image category. To determine if any of these predictors significantly modulated firing rate we generated surrogate-corrected test statistics by shuffling the trial labels 500× to disrupt the relationship between firing rates and image features. Trial labels were only shuffled within category, to ensure the rigorous maintenance of category-specific effects. We then re-fit the model to the shuffled data, and compared the true *F*-value of each predictor to this null distribution. If a predictor exceeded the 95th percentile of the values in the null distribution for that predictor, then it was considered to be significantly tuned to that predictor. To determine whether a significant number of neurons in a region exhibited significant tuning, we used binomial tests evaluated against a chance level of 5%, and Bonferroni-corrected the *P*-values for multiple comparisons across the number of brain regions.

### Identifying spatially modulated cells

We binned the emotion space spanned by the valence and arousal ratings of the image stimuli into 15 x 15 bins. We used this discretized emotion space to compute a firing-rate map for each neuron. To do so, separately for each neuron, we assigned each image to a bin depending on its valence and arousal rating, counted the number of spikes during the presentation of that image, and divided the number of spikes by the amount of time spent in that bin. Because we used the entire encoding period for each image (2 seconds), bins could only be occupied in multiples of 2 seconds. We then smoothed the firing-rate maps using a Gaussian kernel (size = 5 bins, standard deviation = 1). Participants did not sample the entirety of the emotion space (Figs. S7; S8), which would lead to systematic under-estimations of filter coefficients during smoothing if unsampled locations were simply replaced with 0s to enable smoothing, and then replaced with NaN entries after smoothing. As such, it was necessary to correct for unsampled space to avoid under-estimating filter coefficients. To do so, we first estimated the firing-rate map of each neuron by coding unsampled locations with 0s, and smoothing this map with the Gaussian kernel (“uncorrected firing-rate map”). We then constructed a map of equal size, in which unsampled locations were coded with 0s and sampled locations were coded with 1s. We filtered this map with the same Gaussian kernel, which resulted in a map of smoothing coefficients (“coefficient map”). We then divided the uncorrected firing-rate map by the coefficient map to obtain the final firing-rate map that accounted for unsampled locations (“corrected firing-rate map”). This procedure ensured that unsampled locations did not bias the smoothing results.

To quantify the temporal stability of the firing-rate maps, we computed firing-rate maps separately for the first and the second half of the data, separately for each neuron. We then computed the Pearson correlation between these two firing-rate maps to obtain the split-half correlation of all firing-rate maps^13^. We tested whether the split-half correlations were significantly above 0 for grid-like neurons (indicating temporal stability) and whether they were significantly higher for grid-like neurons than for other neurons.

We estimated spatial information^80^ and grid scores^15,41^ for each smoothed firing-rate map. We computed spatial information by summing over the product of the occupancy density and firing rate for each position, multiplied by the log-ratio of the firing rate divided by the overall mean firing rate. This sum was then divided by the mean firing rate, resulting in the spatial information per spike. We computed grid scores by generating spatial autocorrelation matrices from each neurons’ firing-rate map, which was then systematically rotated at various angles. We computed the Pearson product moment correlation coefficient between each rotated autocorrelation matrix and the original autocorrelation matrix. The grid score was then identified as the difference between the minimum correlation coefficient of the autocorrelation matrices shifted by the peak angles (60^◦^and 120^◦^) and the maximum correlation coefficient of the autocorrelation matrices shifted by the trough angles (30^◦^, 90^◦^and 150^◦^)^18^. To test rotational alternatives to 6-fold symmetry, we performed the same procedure for alternative arrangements including 4-, 5-, 7-, 8-, 9-, and 10-fold symmetry, which involved changing the peak and trough angles in the grid-score computation.

Critically, we only used surrogate-corrected spatial information scores and grid scores, as each measure may be biased by low spike counts and thus result in false-positive identifications of spatially-modulated cells^42,81^. We utilized two approaches to do this. In the first approach, we shuffled trial labels so that the arousal and valence values associated with each image were randomly interchanged. In the second approach, we circularly shifted the spike train by different time intervals relative to the behavioral data (minimum shift = 1.25 seconds). Both procedures disrupted the association between firing rates and image characteristics while maintaining the general firing-rate properties of the cells. In both approaches, we performed the shuffling/shifting 500 times, each time recomputing the cells’ firing-rate maps and their corresponding spatial metrics. This formed the basis for an empirically derived null distribution for each cell, allowing us to compute the z-score of each neuron’s spatial information and grid score relative to this distribution. Both approaches yielded highly correlated results (Fig. S6), suggesting that both surrogate methods led to similar statistical thresholds for identifying cells with significant tuning.

Furthermore, for grid-like firing we employed a third surrogate approach in which we only shifted the location of firing fields, as this has been shown to further decrease the risk of false-positive grid-cell identifications in rodent data^81^. To do so, we first identified firing fields by identifying peaks in the firing-rate map (at least 1 standard deviation above the mean firing rate), and then searching for adjacent bins that exceeded 66% of the maximum spatial firing rate, while ensuring that fields had to be separated by at least 2 spatial bins. We then randomly shifted each peak, and iterated through shifting nearby bins by the same displacement vector, in order of proximity to the peak bin. If the displacement vector would place the adjacent bins outside the boundaries of the map or overlaid upon a previously shifted bin, we instead shifted the bin to the nearest available space^81^. This approach also yielded highly correlated results with the bin-shuffle approach (Fig. S6) and, critically, identified the same number of grid-like neurons in the entorhinal cortex (*n* = 14). For the final identification of significant grid-like firing we thus used the bin-shuffle method.

We identified the grid-orientation angle of each grid-like neuron by manually identifying the angular offset (in degrees) between the nearest axis of fields in the spatial autocorrelogram and the horizontal axis^43^. When analyzing the pair-wise angular differences between the grid orientations of different cells, we found that grid orientations of grid-like cells did not appear to be clustered around a particular angle (neither for grid-like cells from the same microwire bundle nor for grid-like cells from different microwire bundles).

### Control analyses using rodent grid-cell data

In order to validate our statistical approach and to ensure that the binned occupancy of emotion space was not resulting in false-positive identifications of grid-like firing, we applied our analysis pipeline to entorhinal cortex neuronal spiking (*n* = 296 neurons) recorded from rodents navigating a square spatial environment^41^. Because these data have been rigorously analyzed in prior publications, they represented a “ground truth” dataset for the identification of grid-cell activity. First, to identify the optimal bin count for estimating our firing-rate maps in emotion space, we applied several different bin counts to our data and used the resulting occupancy maps to mask the rodent firing-rate maps. We then computed the correlation between the grid scores (z-scored relative to surrogate grid scores) estimated using the unmasked rodent data and estimated using the masked rodent data. This showed that a bin size of 15 resulted in the least bias in recapturing “true” grid scores from the unmasked rodent data when using the masked rodent data. Next, in order to determine whether the sparser occupancy of emotion space (relative to more uniform navigational sampling) introduced a bias or resulted in false-positive identifications of grid cells, we masked the rodent data using each participant’s occupancy map and computed the grid score of the resulting firing-rate maps. We classified each resulting grid score as a grid cell by utilizing the same threshold as in the original experiment that produced the dataset (grid score > 0)^41^. We then averaged the classifications (grid cell = 1, non-grid cell = 0) across all the participants’ occupancy maps, and compared this average to the “ground-truth” classification, computing the receiver-operating characteristic (ROC) curve and the corresponding area-under-the-curve (AUC). This showed that our method replicated the classification of grid cells in the rodent dataset with high accuracy, without introducing false-positive identification of grid cells (Fig. S8).

### Measuring theta-phase modulation of neuronal spiking

We estimated the instantaneous phase of the microwire LFPs at low frequencies. First, we removed and interpolated the LFP in the window 2 ms before and 8 ms after each action potential^82^. We also detected and removed interictal discharges by identifying large transients in the 25–80 Hz range that were 3 standard deviations above the mean amplitude and shorter than 200 ms. In line with prior work examining theta modulation of neuronal spiking in humans, we analyzed frequencies ranging from 2–10 Hz^48^. In order to analyze fluctuations in the LFP, we estimated 2–10-Hz phase by first identifying peaks, troughs, and midpoints in a broader (30 Hz) lowpass-filtered LFP, and then linearly interpolating between these points with respect to the (2–10 Hz) band-pass filtered signal to estimate phase in the desired frequency range. This phase-interpolation method has been used previously to effectively estimate theta phase in bats^47^, rodents^83,84^, and humans^48^ as an alternative to using the Hilbert transform to estimate the true peaks and troughs of the signal^84,85^. To ensure that phase estimates were not based on an unreliable low-amplitude signal, we computed the instantaneous power of the LFP and discarded those time-points in which the power fell below a 33rd percentile threshold. To measure the theta-modulation index, we analyzed the spike-phase autocorrelation of the unwrapped spike-phases, as in prior research in rodents^86^, bats^47^, and humans^48^. To do so, we computed the Fourier transform of the spike-phase autocorrelation to yield a power spectrum measuring the frequency of spiking relative to the frequency of the 2–10 Hz filtered signal. A peak in this power spectrum near 1 indicated that the neuron was phase-locked to the ongoing oscillation of interest. Peaks in the power spectrum were compared to the 95th percentile of peaks from the distribution of surrogate spectra computed from shuffled spiking data to assess the significance of theta modulation. Rayleigh statistics were also computed, which showed a strong correlation with modulation indices computed from the spike-phase autocorrelogram for the grid-like neurons in EC (Pearson’s correlation, *r* = 0.69, *P* < 0.001).

## Data Availability

Data are available from the corresponding authors upon reasonable request.

## Code Availability

All custom Python scripts will be publicly available on GitHub upon publication.

## Acknowledgements

We are very grateful to all patients who participated in this study. We thank the clinical team of the Freiburg Epilepsy Center, Freiburg im Breisgau, Germany, for their continuous support; Julia Schipp and Tim Guth for help with implementing the task; Philipp Rebmann for help with data collection; and Joshua Jacobs for helpful comments on the manuscript. L.K. received funding via the German Research Foundation (DFG; KU 4060/1-1; Projektnummer 447634521; KU 4060/2-1; Projektnummer 527084865) and was supported by the return program of the Ministry of Culture and Science of North Rhine-Westphalia. L.K., A.B., and A.S.-B. were supported by the Federal Ministry of Education and Research (BMBF; 01GQ1705A). P.C.R. received research grants from the Fraunhofer Society (Munich, Germany) and from the Else Kröner-Fresenius Foundation (Bad Homburg, Germany).

## Author Contributions

L.K. conceived the study; L.K., P.C.R., A.B., and A.S-B. collected the data; S.E.Q. analyzed the data; S.E.Q. and L.K. interpreted the results; S.E.Q. and L.K. wrote the manuscript; and all authors reviewed the final manuscript.

## Declaration of Interests

The authors declare no competing interests.

---

**Figure S1:**
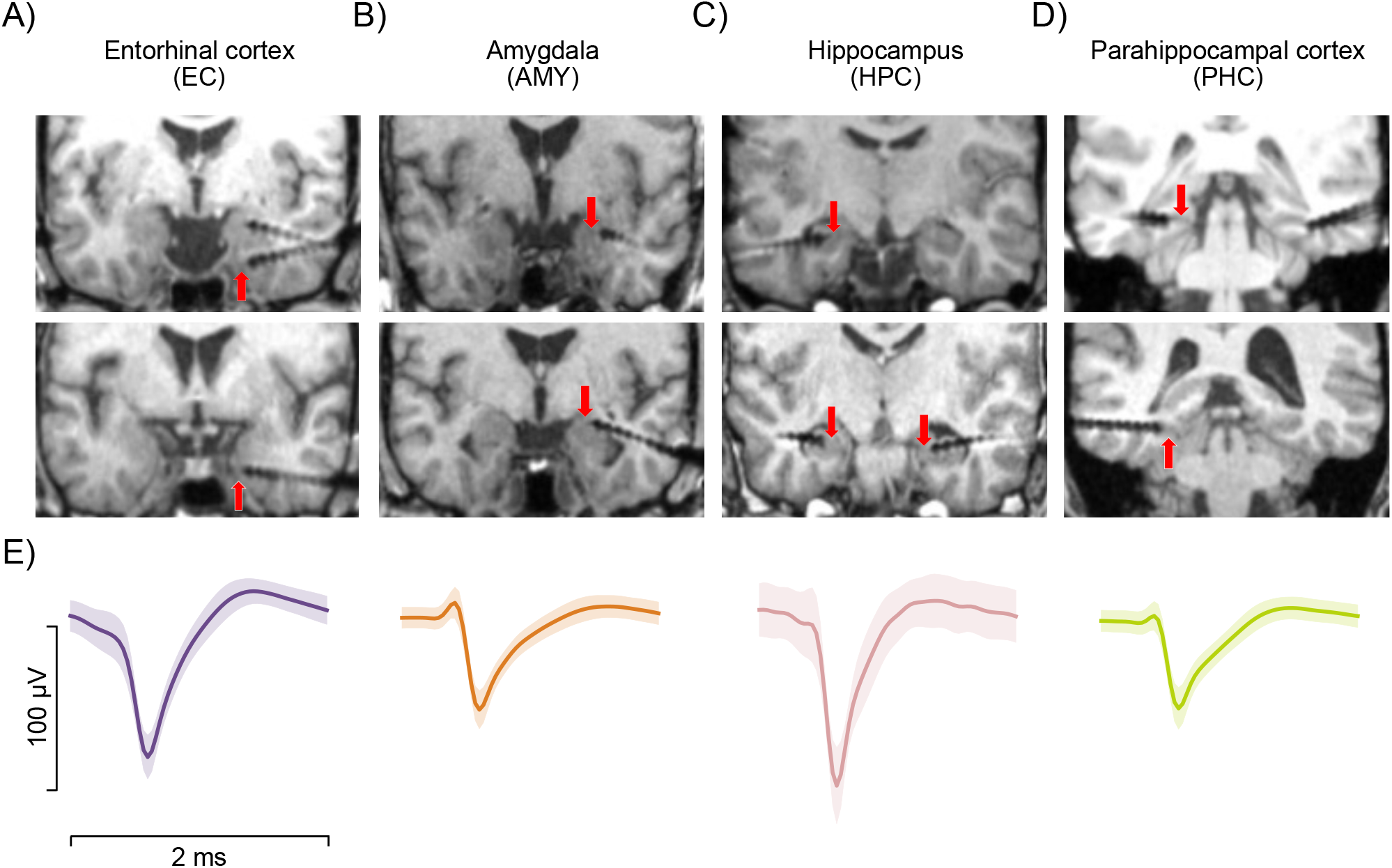
Localization of microwire bundles and example waveforms. (**A**–**D**) Top and middle: example post-operative magnetic resonance imaging (MRI) images indicating the location of microwire bundles in the entorhinal cortex (A), amygdala (B), hippocampus (C), and parahippocampal cortex (D). The putative locations of the microwire bundles are indicated by the red arrows (microwires protrude from the tip of the depth electrodes by 3–5 mm and are often not visible on MRI scans). (**E**) Example average waveforms of putative single units localized to the entorhinal cortex, amygdala, hippocampus, and parahippocampal cortex. Shaded areas denote standard errors.

**Figure S2:**
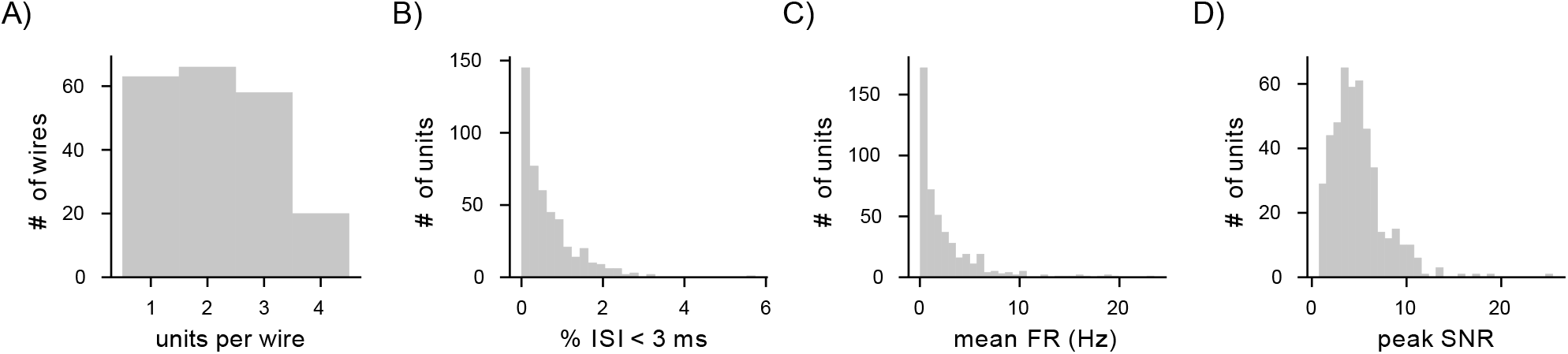
Single-unit quality metrics. (**A**) Histogram of the number of putative single units recorded on each microwire. (**B**) Distribution of the percentages of inter-spike intervals (ISIs) below 3 milliseconds among all ISIs, across single units. (**C**) Average firing rate (FR) across single units (in Hz). (**D**) Peak signal-to-noise ratio (SNR) across single units. #, number.

**Figure S3:**
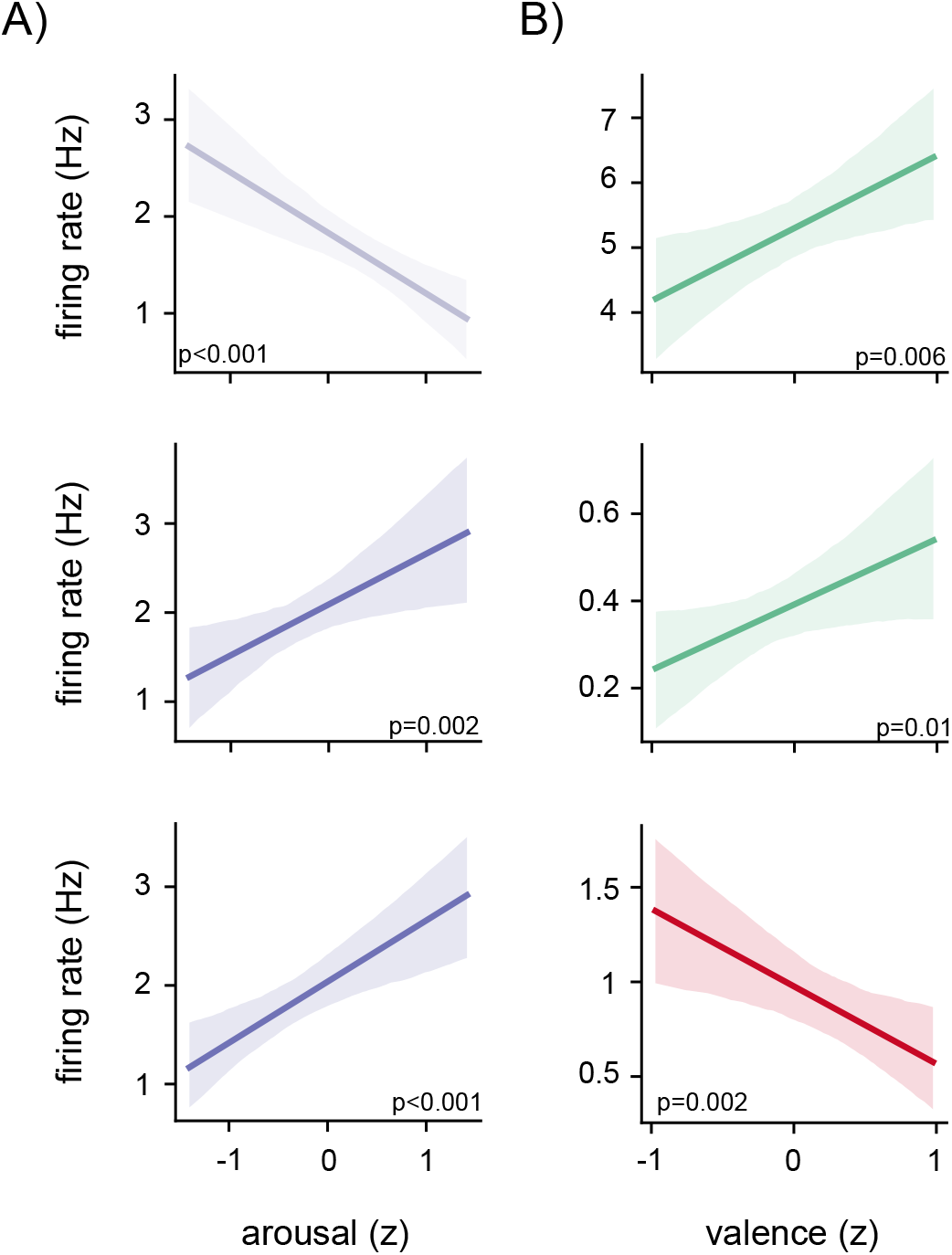
Examples of neurons tuned to arousal and valence alone. (**A**) Linear model fits of the firing rate for example arousal-tuned neurons as a function of z-scored arousal. Shading denotes 95th percentile of bootstrapped model fits. The *P*-value for each neuron is indicated below. (**B**) Linear model fits of the firing rate for example valence-tuned neurons as a function of z-scored valence. Shading denotes 95th percentile of bootstrapped model fits. The *P*-value for each neuron is indicated below.

**Figure S4:**
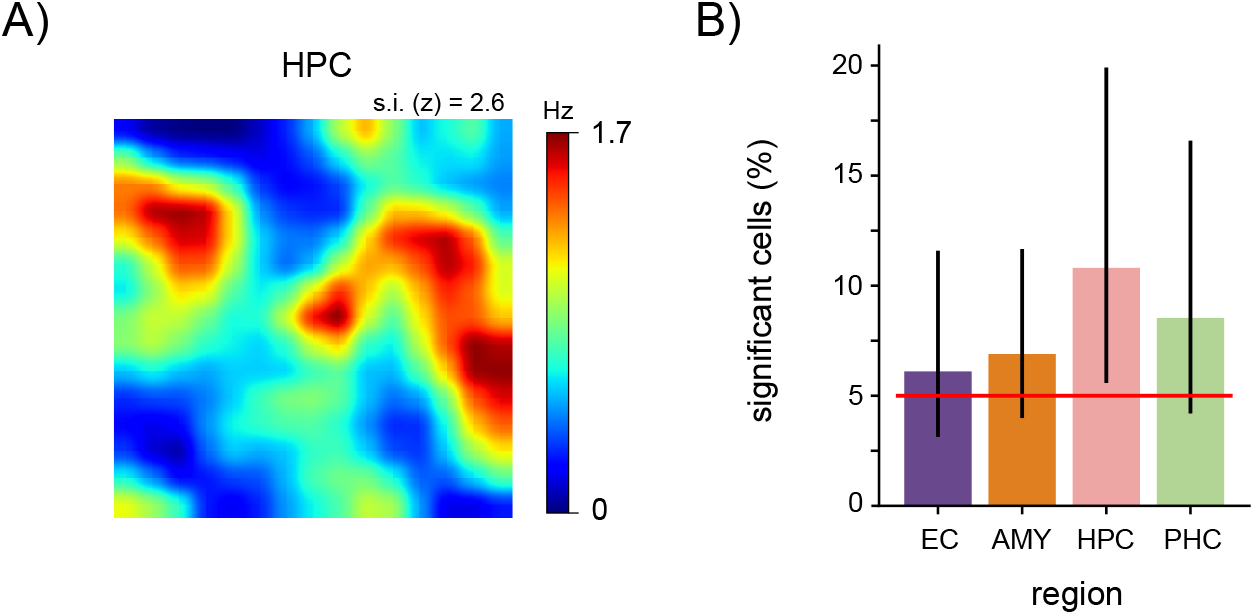
Non-grid spatially tuned neurons. (**A**) Firing-rate map for an example neuron with a significant spatial information score. Warm colors indicate higher firing rates, cool colors indicate lower firing rates. (**B**) Percentage of putative single units in each region that exhibited significant spatial information scores. Vertical lines denote 95% binomial confidence intervals. Red line denotes chance level (5%). None of the regions showed a significant percentage of spatially tuned neurons (binomial tests, all *P*_corr._ < 0.05, Bonferroni corrected for four tests). AMY, amygdala; EC, entorhinal cortex; HPC, hippocampus; PHC, parahippocampal cortex; s.i., spatial information.

**Figure S5:**
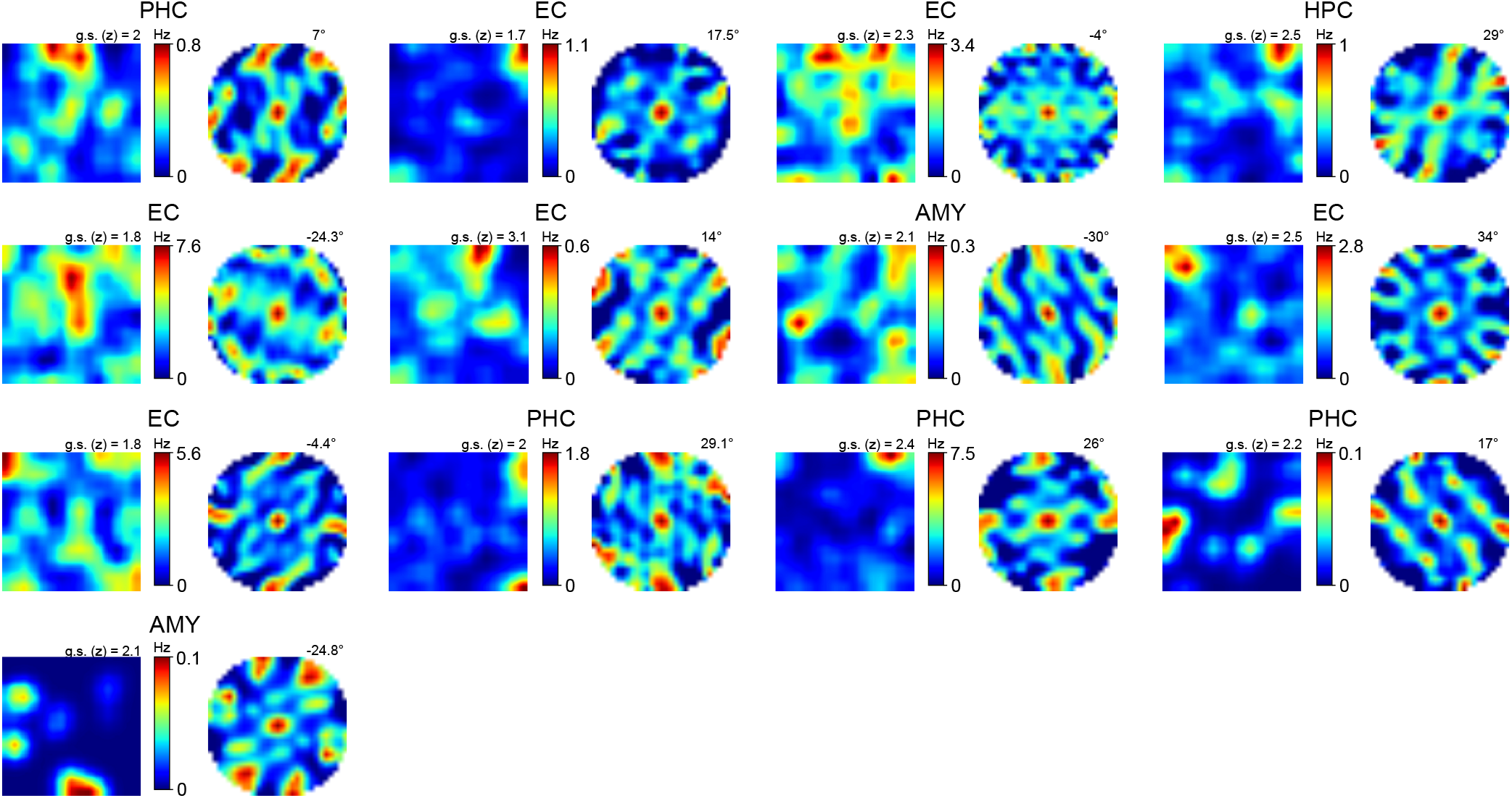
Further examples of grid-like activity in two-dimensional emotion space. For each neuron, firing-rate maps in emotion space are depicted on the left, with colorbar indicating minimum and maximum firing rates. Warm colors denote higher firing rates, cool colors denote lower firing rates. Spatial autocorrelogram is depicted on the right, with the grid-orientation angle stated at the top right. Each neurons’ brain region is indicated above. AMY, amygdala; EC, entorhinal cortex; HPC, hippocampus; PHC, parahippocampal cortex; g.s., grid score.

**Figure S6:**
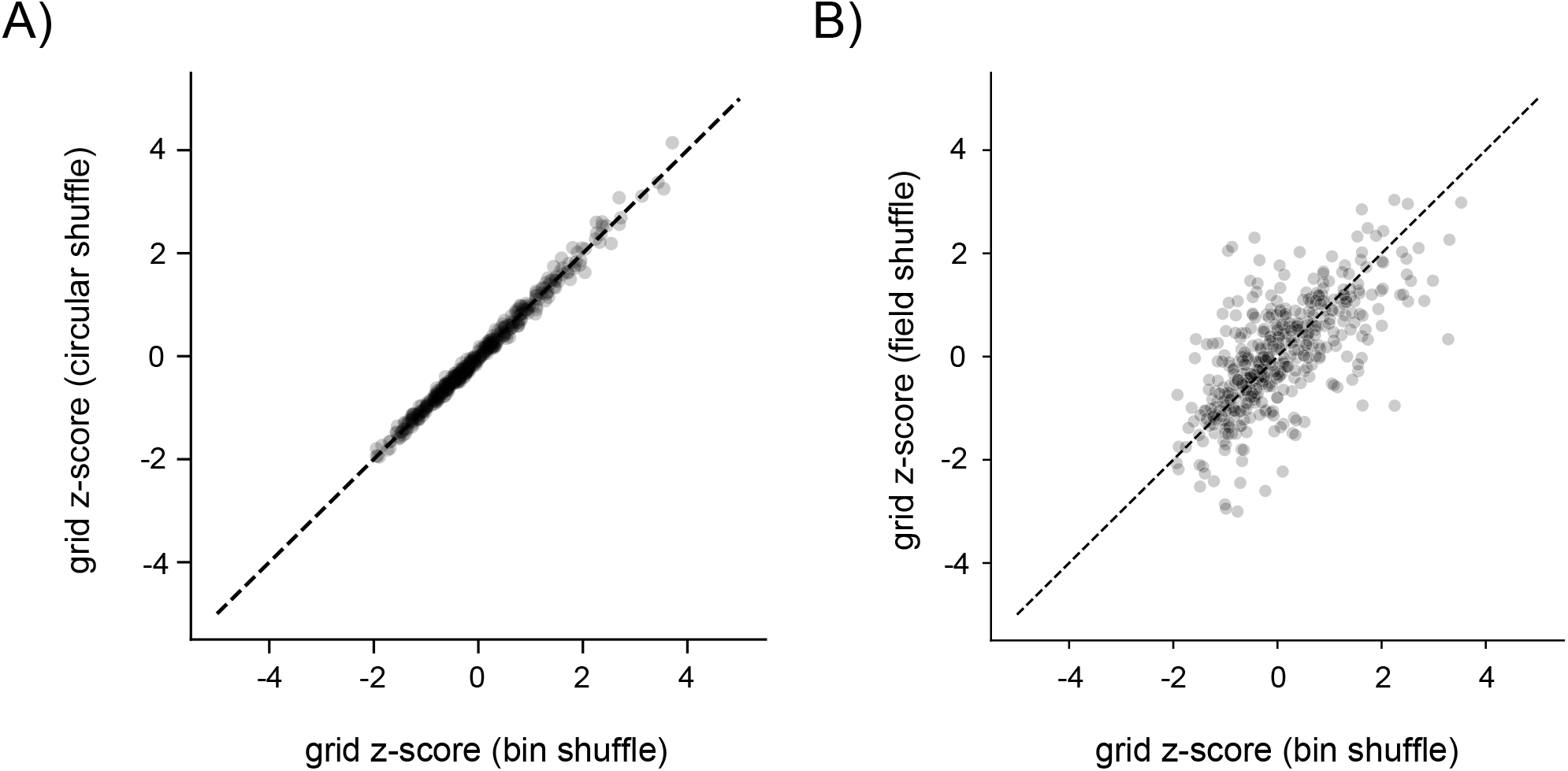
Different techniques for surrogate generation lead to similar z-scored grid scores. (**A**) Correlation between grid scores that were z-scored using the bin-shuffle (x-axis) or the circular-shuffle (y-axis) approach. Dashed line denotes perfect correlation. (**B**) Correlation between grid scores that were z-scored using the bin-shuffle (x-axis) or the field-shuffle (y-axis) approach. Dashed line denotes perfect correlation. We identified 14, 13, and 14 grid-like neurons in the entorhinal cortex using the bin-shuffle, circular-shuffle, and field-shuffle procedure, respectively.

**Figure S7:**
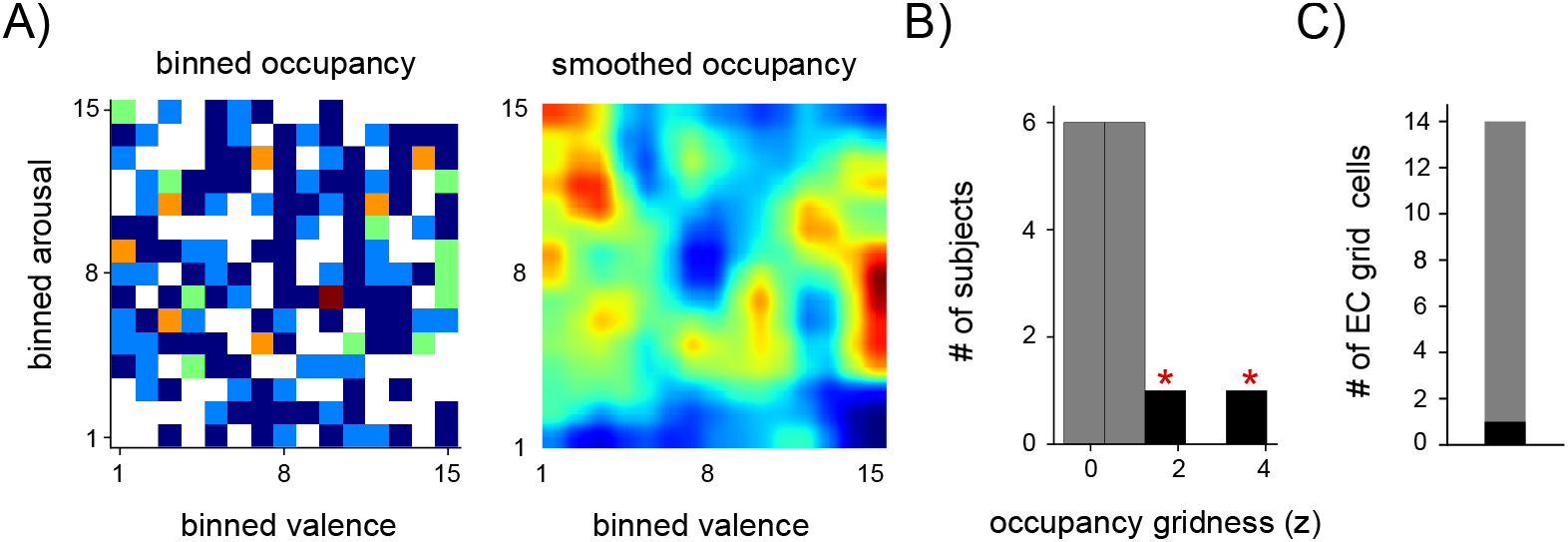
Grid-like behavioral occupancy patterns in two-dimensional emotion space do not account for grid-like neural activity. (**A**) Raw and smoothed occupancy maps for an example participant. Bin-wise occupancy was estimated as the total time the participant viewed images assigned to a particular valence–arousal bin. (**B**) Grid scores computed using the occupancy maps from each session. The occupancy maps of two participants exhibited a significant grid score (black bars with red asterisks). (**C**) Among the 14 entorhinal grid-like cells there was only one cell that was recorded in a participant with a grid-like behavioral occupancy of the emotion-coordinate space. This indicates that the behavioral sampling of emotion space was not a factor that could potentially explain the neurons’ grid-like activity. #, number.

**Figure S8:**
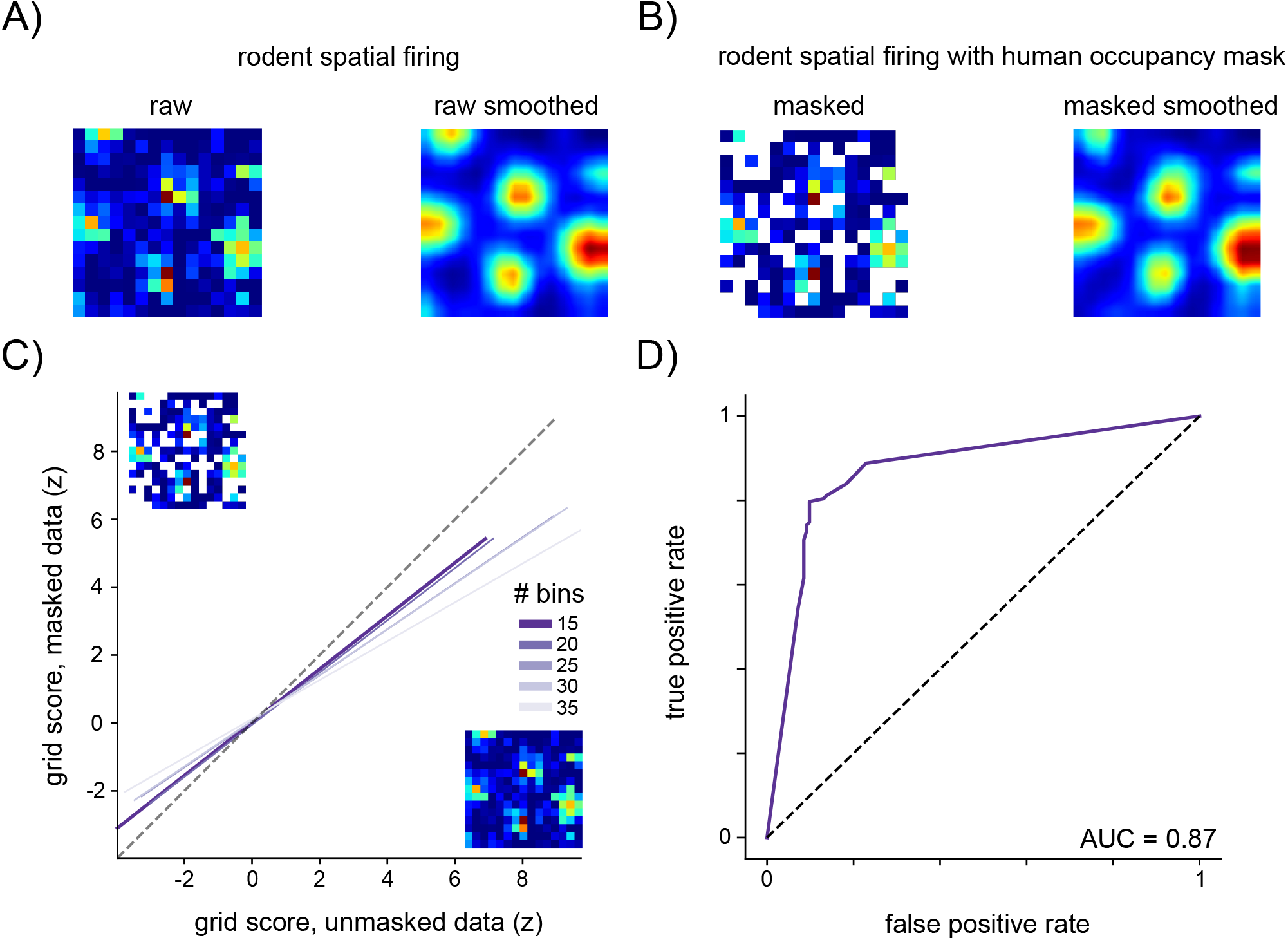
Rationale for using 15 x 15 bins in the analysis of grid-like activity based on an independent rodent grid-cell dataset^41^. (**A**) Firing-rate map of an example grid cell recorded from the entorhinal cortex of a rodent that navigated through a spatial environment. The “raw” and “raw smoothed” firing-rate maps show spiking activity binned according to spatial coordinates using the entire data (i.e., no unvisited locations). (**B**) Left: Raw firing-rate map for the same neuron, masked with the occupancy map from a human participant of our study (unoccupied bins are shown in white). Right: smoothed firing-rate map for the same neuron, based on the masked raw firing-rate map on the left. This procedure simulates the effect of incomplete sampling of emotional space on the detection of grid-like activity in this study. Note the high correspondence between the masked smoothed firing-rate map in B and the raw smoothed firing-rate map in A, indicating that our spatial smoothing procedure adequately dealt with unvisited locations. (**C**) Correlation between grid scores of the raw, smoothed firing-rate maps and their corresponding masked, smoothed firing-rate maps (masked with human occupancy maps), across all neurons from the rodent dataset (*n* = 296). Different colors denote different bin counts, demonstrating that a bin count of 15 x 15 is closest to perfect replication (denoted by the dashed line). This motivated our use of 15 x 15 bins for the analysis of grid-like spiking in emotion space. (**D**) Receiver-operating curve (ROC) for classification of rodent grid cells (*n* = 296) when using a bin count of 15 x 15 bins and masking from the occupancy data of each human participant (purple line). The area under the ROC curve (AUC) is indicated.

**Figure S9:**
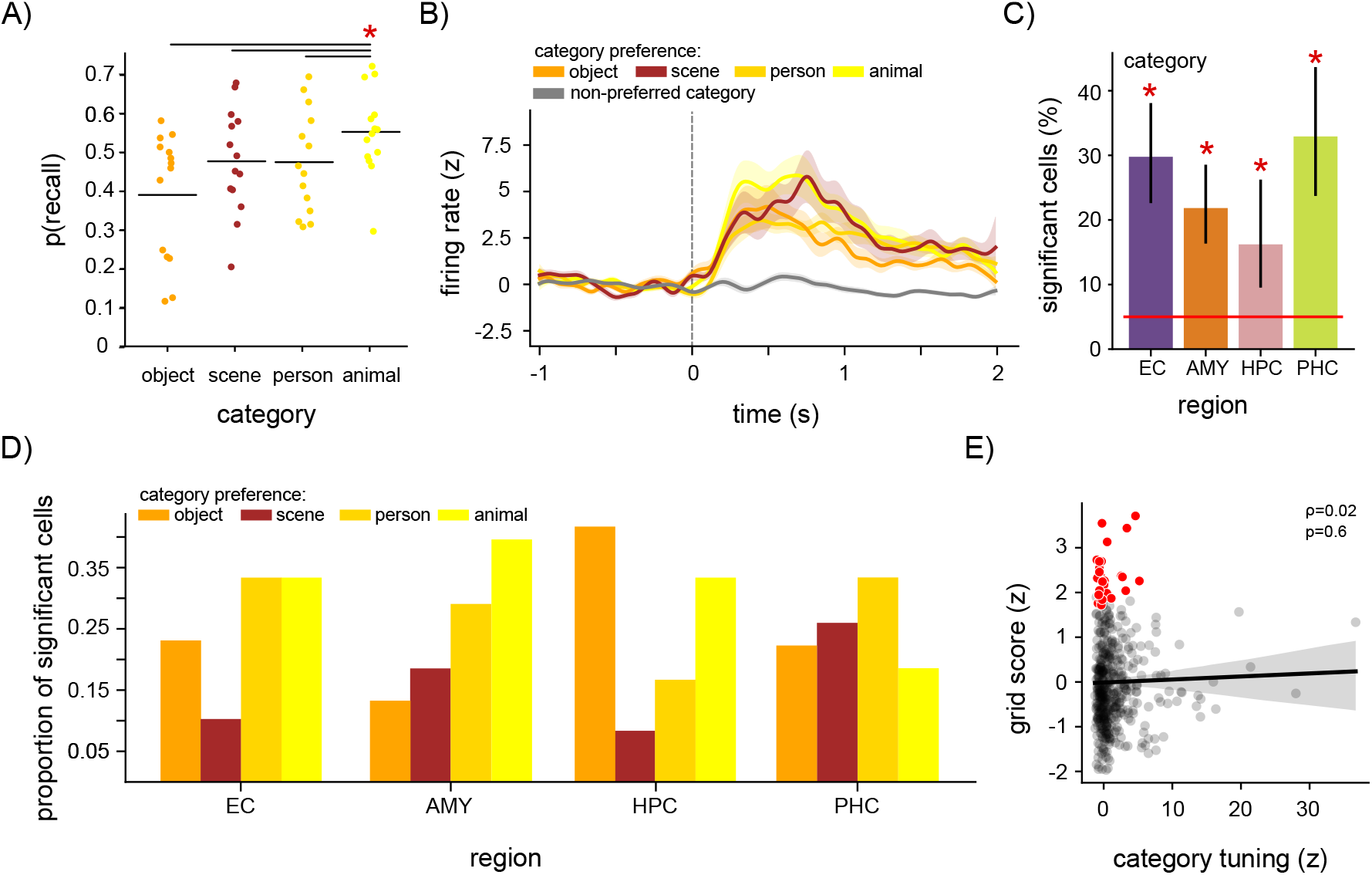
Category tuning does not account for grid-like activity in emotion space. (**A**) Probability of recall across participants (circles) as a function of image category. Horizontal lines denote mean recall probability per image category. Asterisk and horizontal lines at top denote significant *P*-values for coefficients from a logistic regression analysis of recall probability as a function of image category (reference category = animal), indicating that recall probability was higher for animals than for the other categories. (**B**) Mean firing rate (z-scored) of neurons tuned to image category time-locked to image presentation during encoding (dotted line, image onset). Gray line denotes firing rate for the image category with the lowest mean firing rate for each category-tuned neuron. Shaded areas denote the standard error. (**C**) Percentage of putative single units in each region that exhibited significant category tuning. Vertical lines denote 95% binomial confidence intervals. Red line denotes chance level (5%). Asterisks denote significant percentages (binomial tests, *P*_corr._ < 0.05, Bonferroni corrected for four tests). (**D**) Proportion of category-tuned cells split by preferred category and brain region. For example, about 40% of the category-tuned amygdala neurons preferred the image category of animals, in line with previous findings^87^. (**E**) Non-significant correlation between category tuning and emotion grid scores. Line denotes linear regression fit. Shaded areas denote bootstrapped 95% confidence intervals. Red dots denote grid-like neurons across all brain regions. AMY, amygdala; EC, entorhinal cortex; HPC, hippocampus; PHC, parahippocampal cortex.

**Figure S10:**
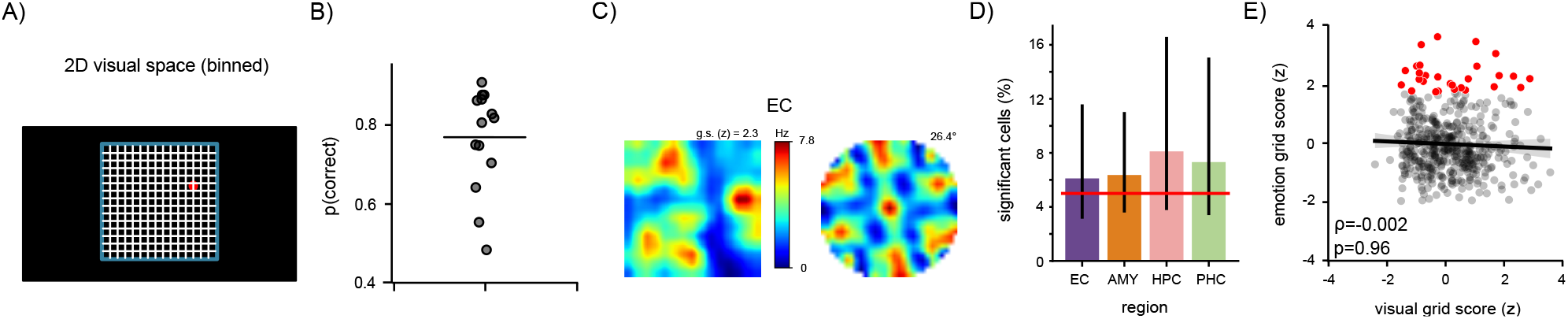
Grid-like tuning to visual space does not account for grid-like activity in emotion space. (**A**) Schematic of the 2D visual space during the distractor task (blue square; 810 x 810 pixels). Black filled rectangle indicates the extent of the laptop screen (1920 x 1080 pixels). During each distractor period, 25 (or 20) red and white dots appeared at random locations in visual space (0.5-s duration per dot), illustrated by the single red dot. Participants were asked to press the space bar whenever they saw a red dot (in order to keep them attentive to the locations of the dots). Participants only saw the red and white dots, but neither the blue square nor the white bins. To test for cells with grid-like tuning in visual space, we estimated firing rates as a function of location in visual space (15 x 15 location bins). As illustrated in Figure 3A, we generated permutation-corrected grid scores relative to empirically generated surrogate null distributions. (**B**) Probability of accurate responses (i.e., pressing the space bar during a red dot) during the distractor period for all participants (one dot per participant). Horizontal line denotes the mean correct response probability of 77%, indicating that participants were attentive to the dot locations in visual space (one-sample *t*-test against 0.5, *P* < 0.001). (**C**) Example of a neuron with grid-like activity in visual space. Firing-rate map in visual space is depicted on the left, with colorbar indicating minimum and maximum firing rates. Warm colors denote higher firing rates, cool colors denote lower firing rates. Spatial autocorrelogram is depicted on the right, with the grid-orientation angle stated at the top right. The neuron’s brain region is indicated above. (**D**) Percentage of putative single units in each region that exhibited significant grid scores in visual space. Vertical lines denote 95% binomial confidence intervals. Red line denotes chance level (5%). We note that we inferred the participants’ viewing locations in visual space from the dot locations, which does not guarantee that participants were indeed looking at these screen locations. Studies with eye tracking are thus better suited to identify grid-like spiking in visual space and may identify higher proportions of visual grid-like neurons (e.g., ref. ^20^). (**E**) Correlation between visual grid scores and emotion grid scores. Line denotes linear regression fit. Shaded areas denote bootstrapped 95% confidence intervals. Number of cells with significant emotion grid scores: 28; number of cells with significant visual grid scores: 31; number of cells with significant emotion and visual grid scores: 3. AMY, amygdala; EC, entorhinal cortex; HPC, hippocampus; PHC, parahippocampal cortex; g.s., grid score.

**Table S1:**
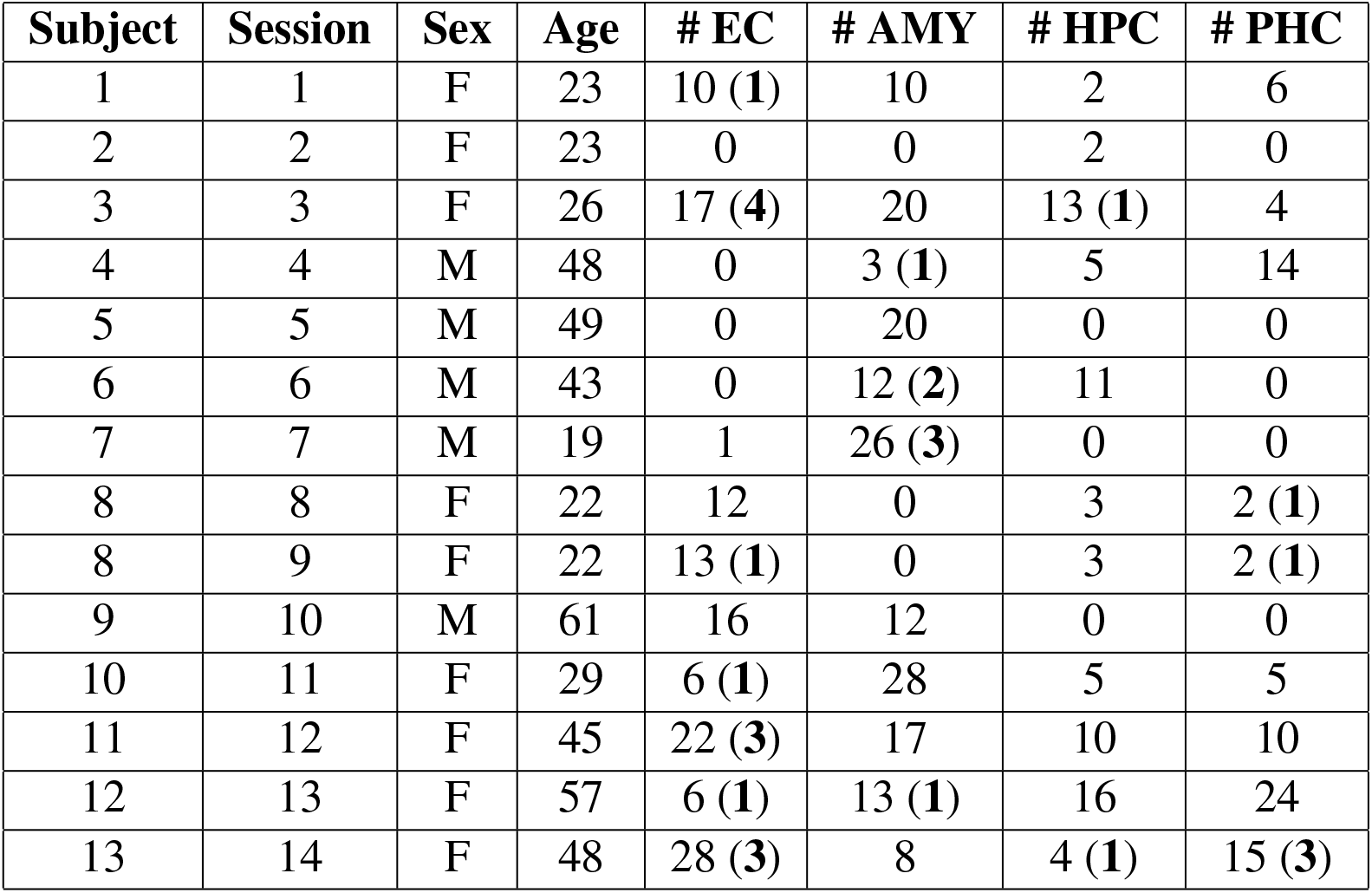
Demographics of participants. Sex, age, and counts of neurons recorded in entorhinal cortex (EC), amygdala (AMY), hippocampus (HPC), and parahippocampal cortex (PHC) for each participant. Grid-like neurons are indicated in parentheses. F, female; M, male; #, number of neurons.

## Notes

### Competing Interest Statement

The authors have declared no competing interest.

