## Supplementary Information for "Neurons in the human entorhinal cortex map abstract emotion space"

| Subject | Session | Sex | Age | # EC | # AMY | # HPC | # PHC |
|---|---|---|---|---|---|---|---|
| 1 | 1 | F | 23 | 10 (**1**) | 10 | 2 | 6 |
| 2 | 2 | F | 23 | 0 | 0 | 2 | 0 |
| 3 | 3 | F | 26 | 17 (**4**) | 20 | 13 (**1**) | 4 |
| 4 | 4 | M | 48 | 0 | 3 (**1**) | 5 | 14 |
| 5 | 5 | M | 49 | 0 | 20 | 0 | 0 |
| 6 | 6 | M | 43 | 0 | 12 (**2**) | 11 | 0 |
| 7 | 7 | M | 19 | 1 | 26 (**3**) | 0 | 0 |
| 8 | 8 | F | 22 | 12 | 0 | 3 | 2 (**1**) |
| 8 | 9 | F | 22 | 13 (**1**) | 0 | 3 | 2 (**1**) |
| 9 | 10 | M | 61 | 16 | 12 | 0 | 0 |
| 10 | 11 | F | 29 | 6 (**1**) | 28 | 5 | 5 |
| 11 | 12 | F | 45 | 22 (**3**) | 17 | 10 | 10 |
| 12 | 13 | F | 57 | 6 (**1**) | 13 (**1**) | 16 | 24 |
| 13 | 14 | F | 48 | 28 (**3**) | 8 | 4 (**1**) | 15 (**3**) |

*Table S1:* **Demographics of participants**. Sex, age, and counts of neurons recorded in entorhinal cortex (EC), amygdala (AMY), hippocampus (HPC), and parahippocampal cortex (PHC) for each participant. Grid-like neurons are indicated in parentheses. F, female; M, male; #, number of neurons.

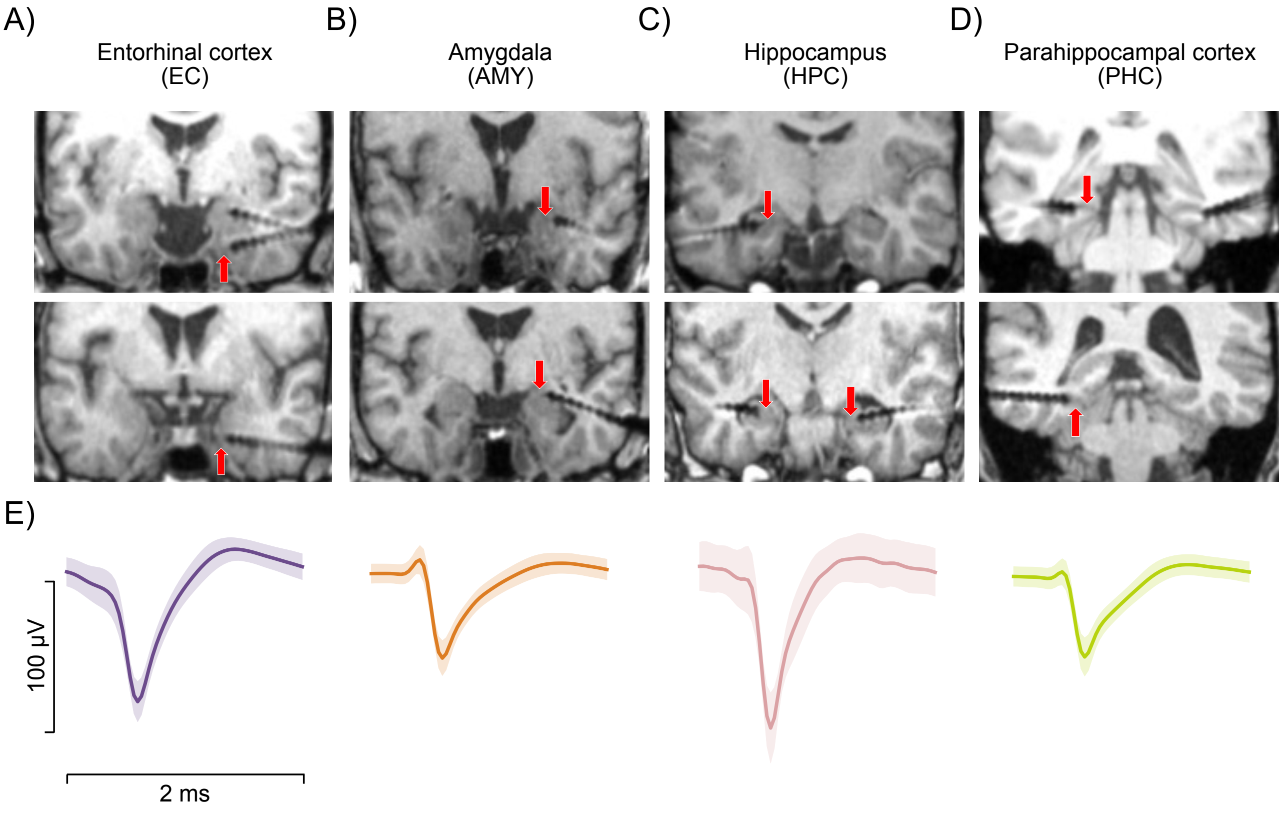


*Figure S1:* **Localization of microwire bundles and example waveforms.** (**A–D**) Top and middle: example post-operative magnetic resonance imaging (MRI) images indicating the location of microwire bundles in the entorhinal cortex (A), amygdala (B), hippocampus (C), and parahippocampal cortex (D). The putative locations of the microwire bundles are indicated by the red arrows (microwires protrude from the tip of the depth electrodes by 3–5 mm and are often not visible on MRI scans). (**E**) Example average waveforms of putative single units localized to the entorhinal cortex, amygdala, hippocampus, and parahippocampal cortex. Shaded areas denote standard errors.

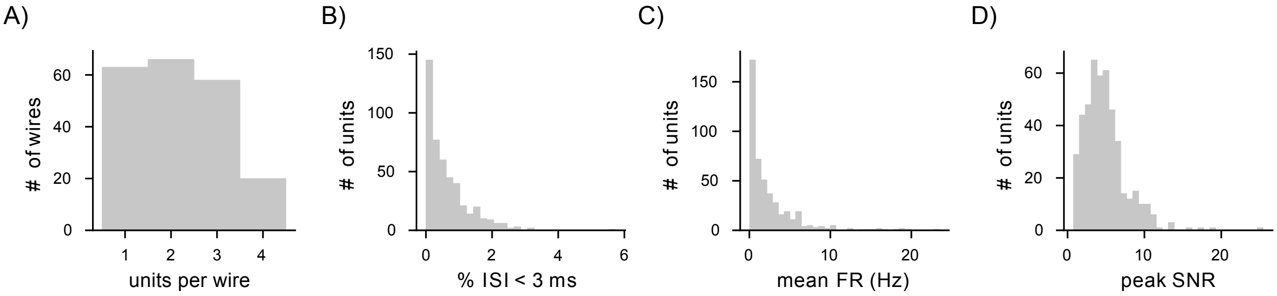


*Figure S2:* **Single-unit quality metrics.** (**A**) Histogram of the number of putative single units recorded on each microwire. (**B**) Distribution of the percentages of inter-spike intervals (ISIs) below 3 milliseconds among all ISIs, across single units. (**C**) Average firing rate (FR) across single units (in Hz). (**D**) Peak signal-to-noise ratio (SNR) across single units. #, number.

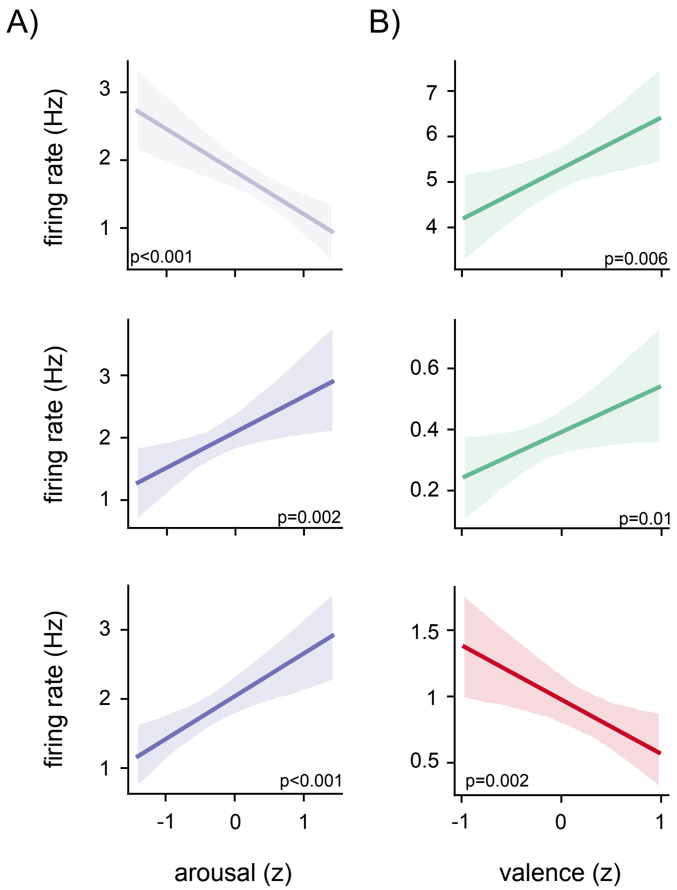


*Figure S3:* **Examples of neurons tuned to arousal and valence alone.** (**A**) Linear model fits of the firing rate for example arousal-tuned neurons as a function of z-scored arousal. Shading denotes 95th percentile of bootstrapped model fits. The *P*-value for each neuron is indicated below. (**B**) Linear model fits of the firing rate for example valence-tuned neurons as a function of z-scored valence. Shading denotes 95th percentile of bootstrapped model fits. The *P*-value for each neuron is indicated below.

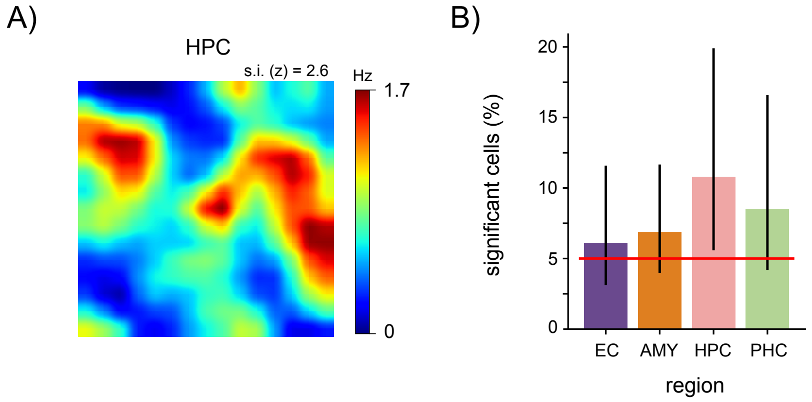


*Figure S4:* **Non-grid spatially tuned neurons.** (**A**) Firing-rate map for an example neuron with a significant spatial information score. Warm colors indicate higher firing rates, cool colors indicate lower firing rates. (**B**) Percentage of putative single units in each region that exhibited significant spatial information scores. Vertical lines denote 95% binomial confidence intervals. Red line denotes chance level (5%). None of the regions showed a significant percentage of spatially tuned neurons (binomial tests, all $P_{\text{corr.}} < 0.05$, Bonferroni corrected for four tests). AMY, amygdala; EC, entorhinal cortex; HPC, hippocampus; PHC, parahippocampal cortex; s.i., spatial information.

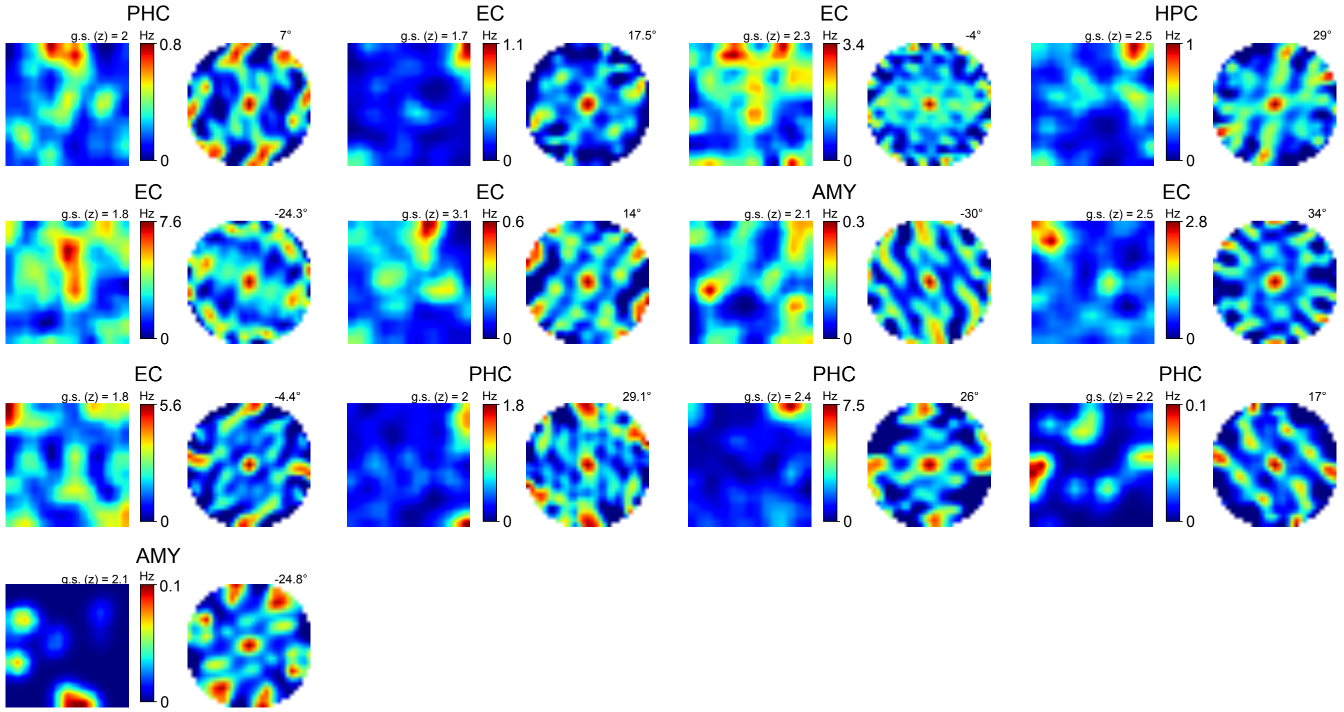


*Figure S5:* **Further examples of grid-like activity in two-dimensional emotion space.** For each neuron, firing-rate maps in emotion space are depicted on the left, with colorbar indicating minimum and maximum firing rates. Warm colors denote higher firing rates, cool colors denote lower firing rates. Spatial autocorrelogram is depicted on the right, with the grid-orientation angle stated at the top right. Each neurons' brain region is indicated above. AMY, amygdala; EC, entorhinal cortex; HPC, hippocampus; PHC, parahippocampal cortex; g.s., grid score.

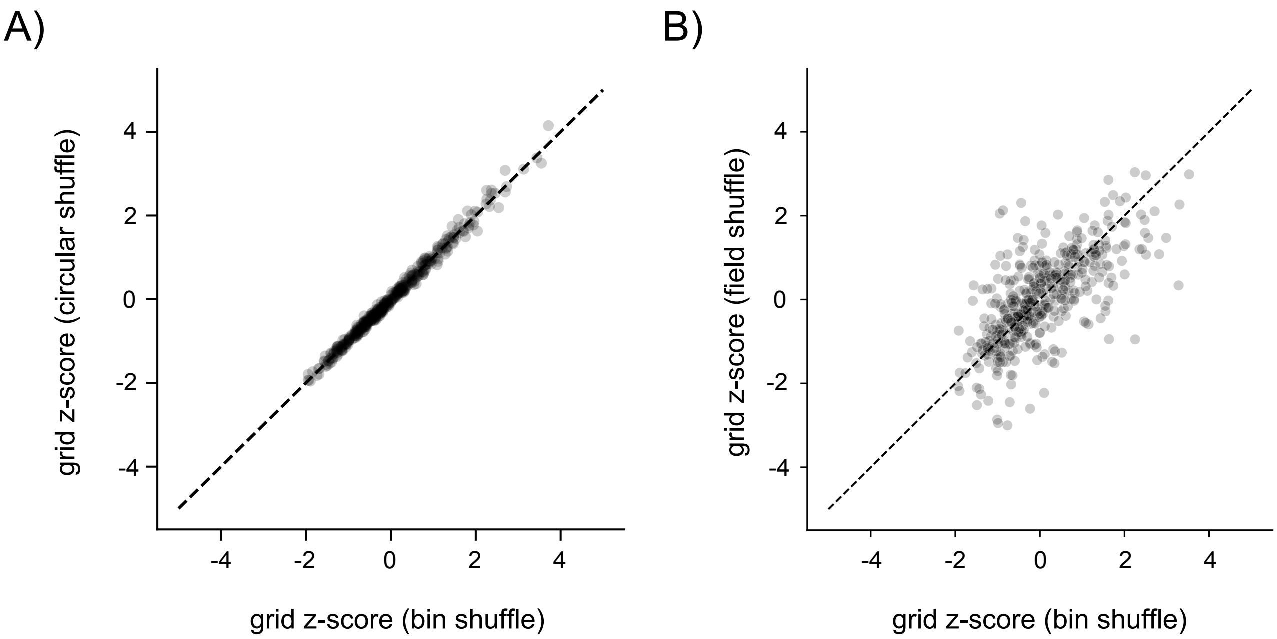


*Figure S6:* **Different techniques for surrogate generation lead to similar z-scored grid scores.** (**A**) Correlation between grid scores that were z-scored using the bin-shuffle (x-axis) or the circular-shuffle (y-axis) approach. Dashed line denotes perfect correlation. (**B**) Correlation between grid scores that were z-scored using the bin-shuffle (x-axis) or the field-shuffle (y-axis) approach. Dashed line denotes perfect correlation. We identified 14, 13, and 14 grid-like neurons in the entorhinal cortex using the bin-shuffle, circular-shuffle, and field-shuffle procedure, respectively.

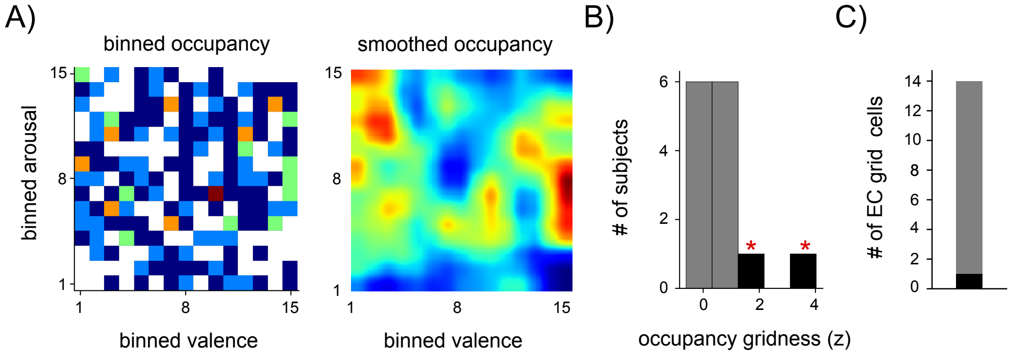


*Figure S7:* **Grid-like behavioral occupancy patterns in two-dimensional emotion space do not account for grid-like neural activity.** (**A**) Raw and smoothed occupancy maps for an example participant. Bin-wise occupancy was estimated as the total time the participant viewed images assigned to a particular valence–arousal bin. (**B**) Grid scores computed using the occupancy maps from each session. The occupancy maps of two participants exhibited a significant grid score (black bars with red asterisks). (**C**) Among the 14 entorhinal grid-like cells there was only one cell that was recorded in a participant with a grid-like behavioral occupancy of the emotion-coordinate space. This indicates that the behavioral sampling of emotion space was not a factor that could potentially explain the neurons' grid-like activity. #, number.

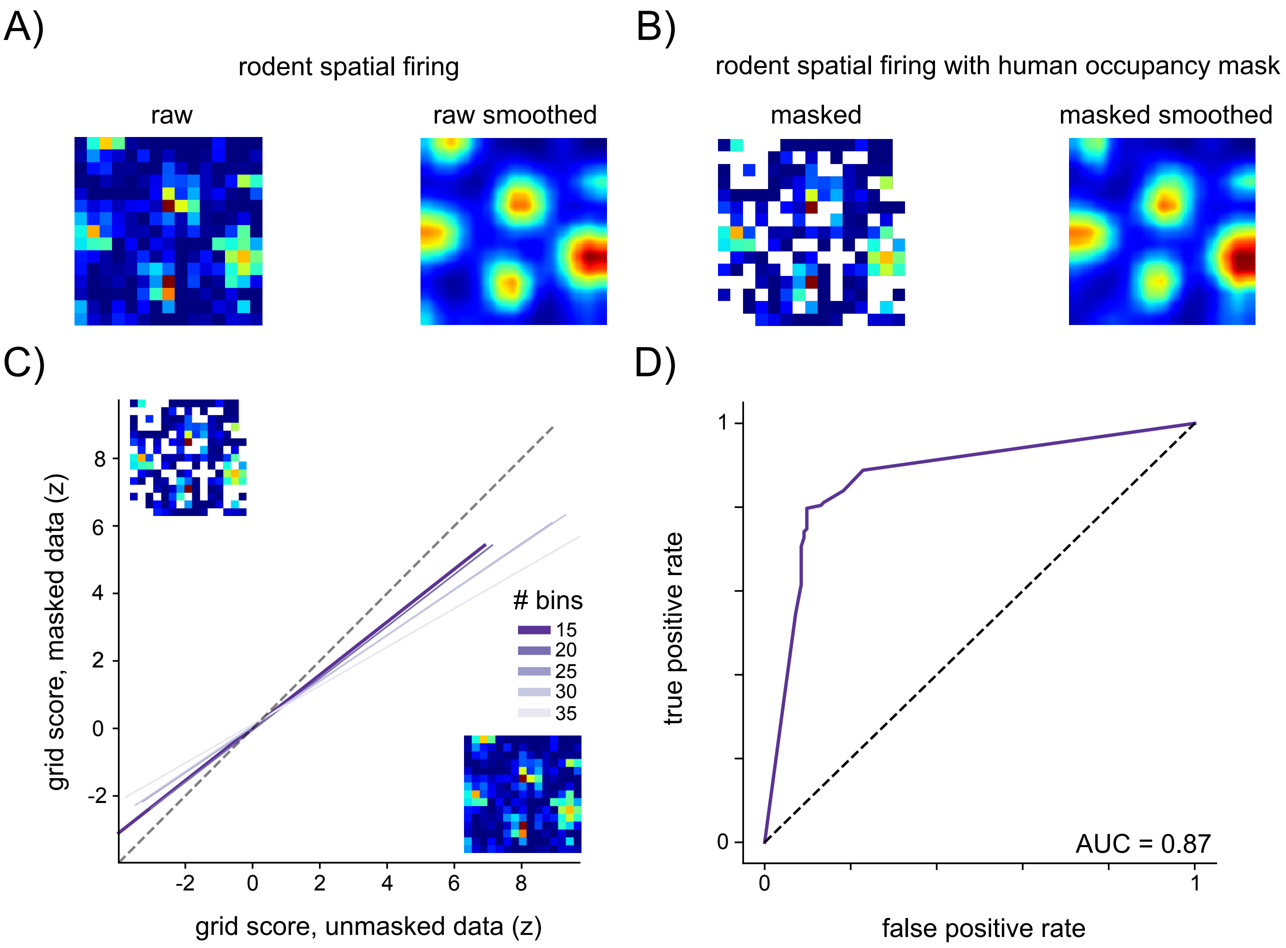


*Figure S8:* **Rationale for using 15 x 15 bins in the analysis of grid-like activity based on an independent rodent grid-cell dataset[41].** (**A**) Firing-rate map of an example grid cell recorded from the entorhinal cortex of a rodent that navigated through a spatial environment. The "raw" and "raw smoothed" firing-rate maps show spiking activity binned according to spatial coordinates using the entire data (i.e., no unvisited locations). (**B**) Left: Raw firing-rate map for the same neuron, masked with the occupancy map from a human participant of our study (unoccupied bins are shown in white). Right: smoothed firing-rate map for the same neuron, based on the masked raw firing-rate map on the left. This procedure simulates the effect of incomplete sampling of emotional space on the detection of grid-like activity in this study. Note the high correspondence between the masked smoothed firing-rate map in B and the raw smoothed firing-rate map in A, indicating that our spatial smoothing procedure adequately dealt with unvisited locations. (**C**) Correlation between grid scores of the raw, smoothed firing-rate maps and their corresponding masked, smoothed firing-rate maps (masked with human occupancy maps), across all neurons from the rodent dataset ($n$ = 296). Different colors denote different bin counts, demonstrating that a bin count of 15 x 15 is closest to perfect replication (denoted by the dashed line). This motivated our use of 15 x 15 bins for the analysis of grid-like spiking in emotion space. (**D**) Receiver-operating curve (ROC) for classification of rodent grid cells ($n$ = 296) when using a bin count of 15 x 15 bins and masking from the occupancy data of each human participant (purple line). The area under the ROC curve (AUC) is indicated.

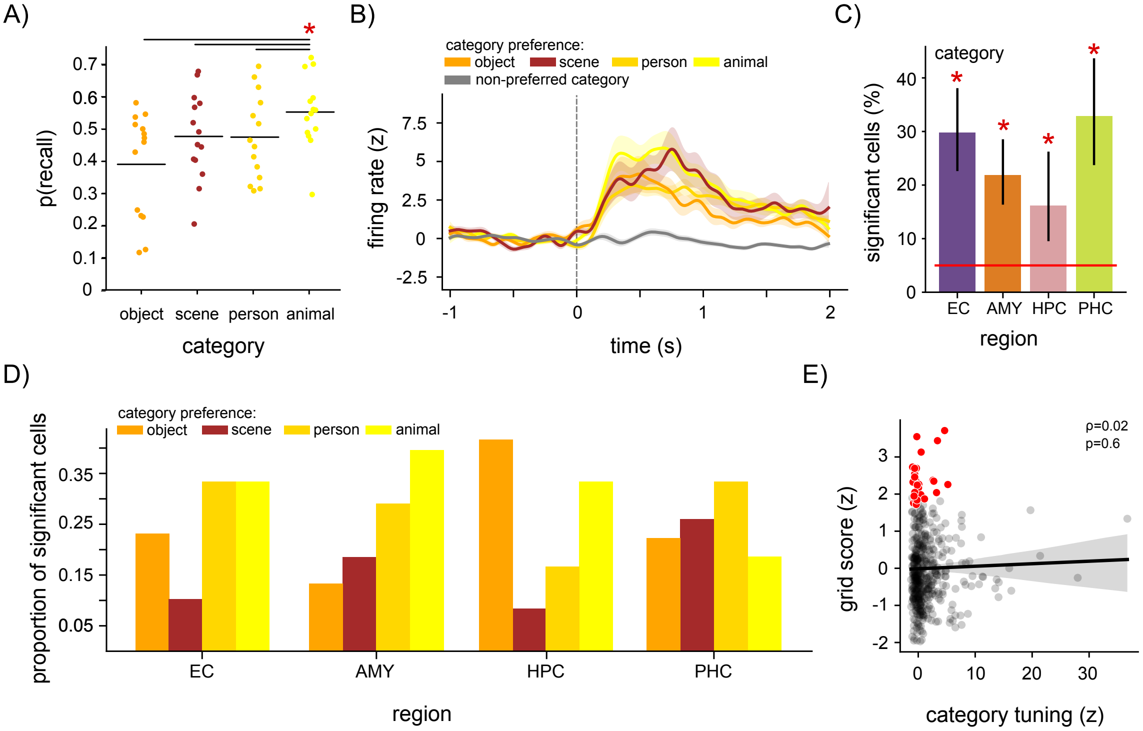


*Figure S9:* **Category tuning does not account for grid-like activity in emotion space.** (**A**) Probability of recall across participants (circles) as a function of image category. Horizontal lines denote mean recall probability per image category. Asterisk and horizontal lines at top denote significant *P*-values for coefficients from a logistic regression analysis of recall probability as a function of image category (reference category = animal), indicating that recall probability was higher for animals than for the other categories. (**B**) Mean firing rate (z-scored) of neurons tuned to image category time-locked to image presentation during encoding (dotted line, image onset). Gray line denotes firing rate for the image category with the lowest mean firing rate for each category-tuned neuron. Shaded areas denote the standard error. (**C**) Percentage of putative single units in each region that exhibited significant category tuning. Vertical lines denote 95% binomial confidence intervals. Red line denotes chance level (5%). Asterisks denote significant percentages (binomial tests, $P_{corr.} < 0.05$, Bonferroni corrected for four tests). (**D**) Proportion of category-tuned cells split by preferred category and brain region. For example, about 40% of the category-tuned amygdala neurons preferred the image category of animals, in line with previous findings[87]. (**E**) Non-significant correlation between category tuning and emotion grid scores. Line denotes linear regression fit. Shaded areas denote bootstrapped 95% confidence intervals. Red dots denote grid-like neurons across all brain regions. AMY, amygdala; EC, entorhinal cortex; HPC, hippocampus; PHC, parahippocampal cortex.

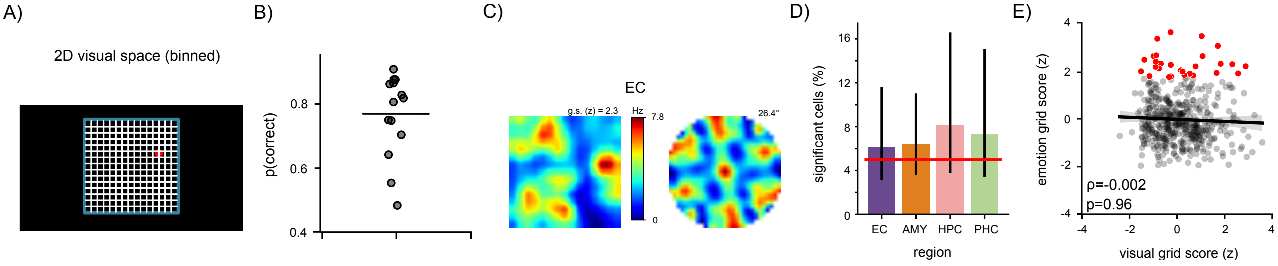


*Figure S10:* **Grid-like tuning to visual space does not account for grid-like activity in emotion space.** (**A**) Schematic of the 2D visual space during the distractor task (blue square; 810 x 810 pixels). Black filled rectangle indicates the extent of the laptop screen (1920 x 1080 pixels). During each distractor period, 25 (or 20) red and white dots appeared at random locations in visual space (0.5-s duration per dot), illustrated by the single red dot. Participants were asked to press the space bar whenever they saw a red dot (in order to keep them attentive to the locations of the dots). Participants only saw the red and white dots, but neither the blue square nor the white bins. To test for cells with grid-like tuning in visual space, we estimated firing rates as a function of location in visual space (15 x 15 location bins). As illustrated in Figure 3A, we generated permutation-corrected grid scores relative to empirically generated surrogate null distributions. (**B**) Probability of accurate responses (i.e., pressing the space bar during a red dot) during the distractor period for all participants (one dot per participant). Horizontal line denotes the mean correct response probability of 77%, indicating that participants were attentive to the dot locations in visual space (one-sample $t$-test against 0.5, $P < 0.001$). (**C**) Example of a neuron with grid-like activity in visual space. Firing-rate map in visual space is depicted on the left, with colorbar indicating minimum and maximum firing rates. Warm colors denote higher firing rates, cool colors denote lower firing rates. Spatial autocorrelogram is depicted on the right, with the grid-orientation angle stated at the top right. The neuron's brain region is indicated above. (**D**) Percentage of putative single units in each region that exhibited significant grid scores in visual space. Vertical lines denote 95% binomial confidence intervals. Red line denotes chance level (5%). We note that we inferred the participants' viewing locations in visual space from the dot locations, which does not guarantee that participants were indeed looking at these screen locations. Studies with eye tracking are thus better suited to identify grid-like spiking in visual space and may identify higher proportions of visual grid-like neurons (e.g., ref. [20]). (**E**) Correlation between visual grid scores and emotion grid scores. Line denotes linear regression fit. Shaded areas denote bootstrapped 95% confidence intervals. Number of cells with significant emotion grid scores: 28; number of cells with significant visual grid scores: 31; number of cells with significant emotion and visual grid scores: 3. AMY, amygdala; EC, entorhinal cortex; HPC, hippocampus; PHC, parahippocampal cortex; g.s., grid score.
